# RoboCOP: Jointly computing chromatin occupancy profiles for numerous factors from chromatin accessibility data

**DOI:** 10.1101/2020.06.03.132001

**Authors:** Sneha Mitra, Jianling Zhong, David M. MacAlpine, Alexander J. Hartemink

## Abstract

Chromatin is the tightly packaged structure of DNA and protein within the nucleus of a cell. The arrangement of different protein complexes along the DNA modulates and is modulated by gene expression. Measuring the binding locations and level of occupancy of different transcription factors (TFs) and nucleosomes is therefore crucial to understanding gene regulation. Antibody-based methods for assaying chromatin occupancy are capable of identifying the binding sites of specific DNA binding factors, but only one factor at a time. On the other hand, epigenomic accessibility data like ATAC-seq, DNase-seq, and MNase-seq provide insight into the chromatin landscape of all factors bound along the genome, but with minimal insight into the identities of those factors. Here, we present RoboCOP, a multivariate state space model that integrates chromatin information from epigenomic accessibility data with nucleotide sequence to compute genome-wide probabilistic scores of nucleosome and TF occupancy, for hundreds of different factors at once. We apply RoboCOP to MNase-seq data to elucidate the protein-binding landscape of nucleosomes and 150 TFs across the yeast genome. Using available protein-binding datasets from the literature, we show that our model predicts the binding of these factors genome-wide more accurately than existing methods.

## 1 INTRODUCTION

A cell’s chromatin consists of the genome and all the proteins and protein complexes arrayed along it. The arrangement of proteins along the genome determines whether and to what extent the cell’s various genes are expressed. Therefore, deciphering the chromatin landscape— the positions of all the different proteins bound to the DNA—is crucial to developing a more mechanistic and predictive understanding of gene regulation.

Two important types of DNA binding factors (DBFs) are transcription factors (TFs) and nucleosomes. TFs are gene regulatory proteins that activate or repress the transcription of genes by binding with specific sequence preferences to sites along the DNA. Nucleosomes form when 147 base pairs of DNA are wrapped around an octamer of histone proteins. They have lower sequence specificity than TFs, but still exhibit a preference for a periodic arrangement of dinucleotides that facilitates DNA wrapping (1). Likened to beads on a string, nucleosomes are positioned fairly regularly along the DNA, occupying about 81% of the genome in the case of the yeast *Saccharomyces cerevisiae* (2). In taking up their respective positions, nucleosomes contribute to the regulation of gene expression in part by allowing or blocking TFs from occupying their putative binding sites. Useful models of the chromatin landscape must therefore be able to simultaneously represent and reason about many DBFs at once, and must explicitly account for the way they compete with one another to bind the genome.

The binding locations of DBFs have been assayed extensively at high resolution with antibody-based methods (3; 4; 5). However, these methods are limited to assaying only one particular factor at a time, and require a separate antibody for each factor. Consequently, using this approach to identify the binding locations of myriad different DBFs is extremely expensive and laborious, especially if we are interested to study how the chromatin landscape changes dynamically across time or in response to changing environmental conditions. In such scenarios, antibody-based methods are often used to assay a small number of important histone modifications, and then computational algorithms integrate the multiple datasets to infer broad segments of ‘epigenomic states’ that can then be associated with larger regulatory loci like promoters and enhancers (6; 7; 8; 9).

In contrast to antibody-based methods, chromatin accessibility assays probe unoccupied, or open, regions of the chromatin, thereby telling us indirectly about the genomic regions occupied by all the bound proteins. Chromatin accessibility data can be generated in a few different ways, including transposon insertion (ATAC-seq), enzymatic cleavage (DNase-seq), or enzymatic digestion (MNase-seq). In the latter, the endo-exonuclease MNase is used to digest unbound DNA, leaving behind undigested fragments of bound DNA. Paired-end sequencing of these fragments reveals not only their location but also their length, yielding information about the length of protein-bound sites throughout the genome. MNase-seq has been widely used to study nucleosome positions (10; 11), but evidence of TF binding sites has also been observed in the data (12; 13). We set out to explore whether a sufficiently sophisticated computational model might be able to leverage this kind of data to identify the precise binding locations of numerous different DBFs at once.

In earlier work, we developed COMPETE to compute a probabilistic landscape of DBF occupancy along the genome (14). COMPETE considers DBFs binding to the genome from the perspective of a thermodynamic ensemble, where the DBFs are in continual competition to occupy locations along the genome and their chances of binding are affected by their concentrations, akin to a repeated game of ‘musical chairs’. COMPETE output depends only on genome sequence (which is static) and DBF concentrations (which may be dynamic); it makes no use of experimental data so its predictions of the chromatin landscape are entirely theoretical. We later developed a modified version of COMPETE to estimate DBF concentrations by maximizing the correlation between the output of COMPETE’s theoretical model and an MNase-seq signal, improving the reported binding landscape (15). However, this modified version still does not incorporate chromatin accessibility data directly into the underlying probabilistic model.

Here, we present RoboCOP (**robo**tic **c**hromatin **o**ccupancy **p**rofiler), a new method that integrates chromatin accessibility data and genomic sequence to produce accurate chromatin occupancy profiles of the genome. With nucleotide sequence and chromatin accessibility data as input, RoboCOP uses a multivariate hidden Markov model (HMM) (16) to compute a probabilistic occupancy landscape of hundreds of DBFs genome-wide at single-nucleotide resolution. In this paper, we use paired-end MNase-seq data to predict TF binding sites and nucleosome positions across the entire *Saccharomyces cerevisiae* genome. We validate our nucleosome positioning predictions using high-precision annotations resulting from a chemical cleavage method (17), and our TF binding site predictions using annotations reported by ChIP-chip (18), ChIPexo (5), and ORGANIC (19) experiments. RoboCOP is the first method to elucidate the chromatin landscape of the genome from MNase-seq data, and can be used to study how chromatin responds dynamically to changing environmental conditions.

## 2 MATERIALS AND METHODS

### 2.1 MNase-seq fragments of different lengths provide information about different kinds of DNA binding factors

Our high-resolution paired-end MNase-seq data can be plotted in two dimensions by representing every fragment as a point, whose *x*-coordinate is the genomic location of its midpoint and whose *y*-coordinate is its length, thereby capturing both the fragment length and location distributions at single base pair resolution. As can be seen from the region in Fig. 1a, gene bodies mostly contain fragments about 150 bases long, corresponding to nucleosomes. Promoter regions contain shorter fragments, often associated with TF binding sites. Each of the two promoter regions in Fig. 1a has an annotated Abf1 binding site (18) that can explain the enrichment of short fragments nearby.

**Figure 1.**
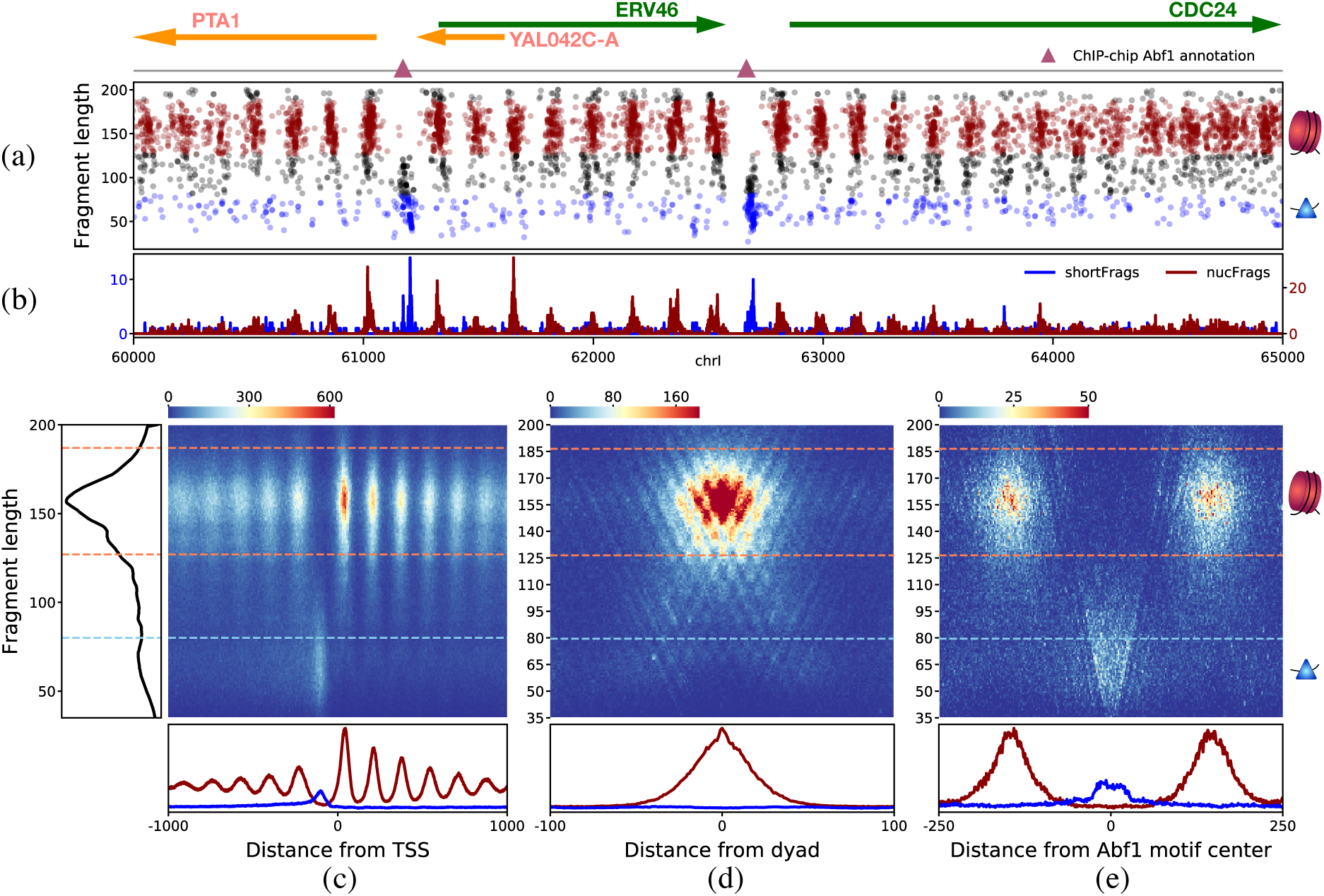
Paired-end MNase-seq data is informative about the binding of both nucleosomes and smaller DBFs, such as transcription factors. (a) Two-dimensional plot of MNase-seq fragments from positions 60,000 to 65,000 of yeast chromosome I. Each fragment is plotted based on its length (*y*-axis) and the genomic location of its midpoint (*x*-axis). Nucleosome-sized fragments (`nucFrags`, length 157±30) are colored red, while shorter fragments corresponding to smaller proteins (`shortFrags`, length ≤80) are colored blue. Above the plot are genomic annotations for this region, with Watson strand genes depicted as green arrows and Crick strand genes as orange. Below the gene annotations, known TF binding sites (18) are indicated using triangles. This region contains two annotated binding sites for Abf1 (pink). (b) Aggregate numbers of red and blue dots at each genomic position in (a), resulting in the one-dimensional `nucFrags` and `shortFrags` signals, respectively. (c) Composite heatmap of MNase-seq fragments around all yeast genes, centered on each gene’s TSS. Panels along the side and bottom show marginal densities. The side panel shows that `nucFrags` predominate, consistent with the fact that over 80% of the yeast genome is occupied by nucleosomes (2), but the bottom panel clarifies that `nucFrags` and `shortFrags` are positioned differently with respect to genes. `nucFrags` appear in tandem arrays within gene bodies, with particularly strong enrichment (deep red) at +1 nucleosomes just downstream of the TSS. In contrast, `shortFrags` are enriched in the nucleosome-free promoter region just upstream of the TSS. (d) Composite heatmap of MNase-seq fragments around the 2000 most well-positioned nucleosomes in the yeast genome (17), centered on each nucleosome’s dyad. The `nucFrags` signal peaks precisely at the dyad and decreases symmetrically in either direction. (e) Composite heatmap of MNase-seq fragments around all annotated Abf1 binding sites (18) in the yeast genome, centered on each site’s motif. Note the clear enrichment of `shortFrags` near Abf1 sites.

Since the degree of MNase digestion can influence the fragment length distribution (20; 21), we plotted the MNase-seq fragments around transcription start sites (TSSs) to get an estimate of the length of the fragments corresponding to nucleosomes and TF binding sites (Fig. 1c). We find that fragments of length 157 have the highest frequency (left panel in Fig. 1c). Given that nucleosomes are about this size and occupy about 81% of the yeast genome (2), fragments of this length generally correspond to nucleosomes. We denote all fragments whose length is 157±30 to be nucleosomal fragments, or `nucFrags` for short. The midpoints of `nucFrags` are depicted in red dots in Fig. 1a. As expected, these `nucFrags` occur in tandem arrays within gene bodies but are generally absent from promoters (Figs. 1a,c). Fragments are particularly concentrated at the +1 nucleosome position in Fig. 1c, just downstream of the TSS, because the +1 nucleosome is usually well-positioned. Furthermore, the marginal density of the midpoints of these fragments around annotated nucleosome dyads (17) peaks precisely at the dyad, with counts dropping nearly symmetrically in either direction (Fig. 1d). This makes sense because MNase digests linker regions, leaving behind undigested DNA fragments wrapped around histone octamers. So the midpoint counts of these `nucFrags` would be highest at the annotated dyads and decrease on moving away from the dyad.

In addition, it has been shown that shorter fragments in MNase-seq provide information about TF binding sites (12). To verify that we see this signal in our data, both the composite plot in Fig. 1c and the genomic region in Fig. 1a reveal that promoter regions are enriched with shorter fragments. The promoter region is often bound by specific and general TFs that aid in the transcription of genes. To ensure that the MNase-seq signal in these promoter regions is not just noise, we plot the MNase-seq midpoints around a set of annotated TF binding sites (Fig. 1e). We choose the well-studied TF, Abf1, because it has multiple annotated binding sites across the genome. On plotting the MNase-seq midpoint counts around these annotated binding sites, we notice a clear enrichment of short fragments at the binding sites. We denote these short fragments of length less than 80 as `shortFrags`. The midpoints of `shortFrags` are plotted as blue dots in Fig. 1a. Unlike the midpoint counts of the `nucFrags` which have a symmetrically decreasing shape around the nucleosome dyads (Fig. 1d), the midpoint counts of `shortFrags` are more uniformly distributed within the binding site (Fig. 1e). The `shortFrags` signal at the Abf1 binding sites is noisier than the MNase signal associated with nucleosomes. One reason for this increased noise is that fragments protected from digestion by bound TFs may be quite small, and the smallest fragments (of length less than 27 in our case) are not even present in the dataset due to sequencing and alignment limitations.

We ignore fragments of intermediate length (81–126) in our analysis, though these could provide information about other kinds of complexes along the genome, like hexasomes (22). Such factors would also be important for a complete understanding of the chromatin landscape, but we limit our analysis here to studying the occupancy of nucleosomes and TFs. For the subsequent sections of this paper, we only consider the midpoint counts of `nucFrags` and `shortFrags` as depicted in red and blue respectively in Fig. 1a. We further simplify the two-dimensional plot in Fig. 1a to form two one-dimensional signals by separately aggregating the midpoint counts of `nucFrags` and `shortFrags`, as shown in Fig. 1b.

### 2.2 RoboCOP model structure and transition probabilities

RoboCOP is a multivariate hidden Markov model (HMM) for jointly computing genome-wide chromatin occupancy profiles using nucleotide sequence and chromatin accessibility data as observables (Fig. 2). The HMM structure has been adapted from (14). Let the number of TFs be *K*. Let ***π***_1_, …, ***π***_*K*_ denote the models for the *K* TFs, and let ***π***_*K*+1_ denote the model for nucleosomes. To simplify notation, we consider an unbound DNA nucleotide to be occupied by a special ‘empty’ DBF (15); suggestively, let ***π***_0_ denote this model. Therefore, we have *K* + 2 DBF models in total, and we use a central non-emitting (‘silent’) state to simplify transitions among them. The HMM may transition from this central silent state to any one of the *K* + 2 DBF models; at the end of each DBF model, the HMM always transitions back to the central silent state (Fig. 2b, Fig. S1). This approach assumes DBFs bind independently of their neighbors, and each DBF therefore has just a single transition probability associated with it. The transition probabilities from the central state to the various DBFs are denoted as {*α*_0_, …, *α*_*K*+1_}.

**Figure 2.**
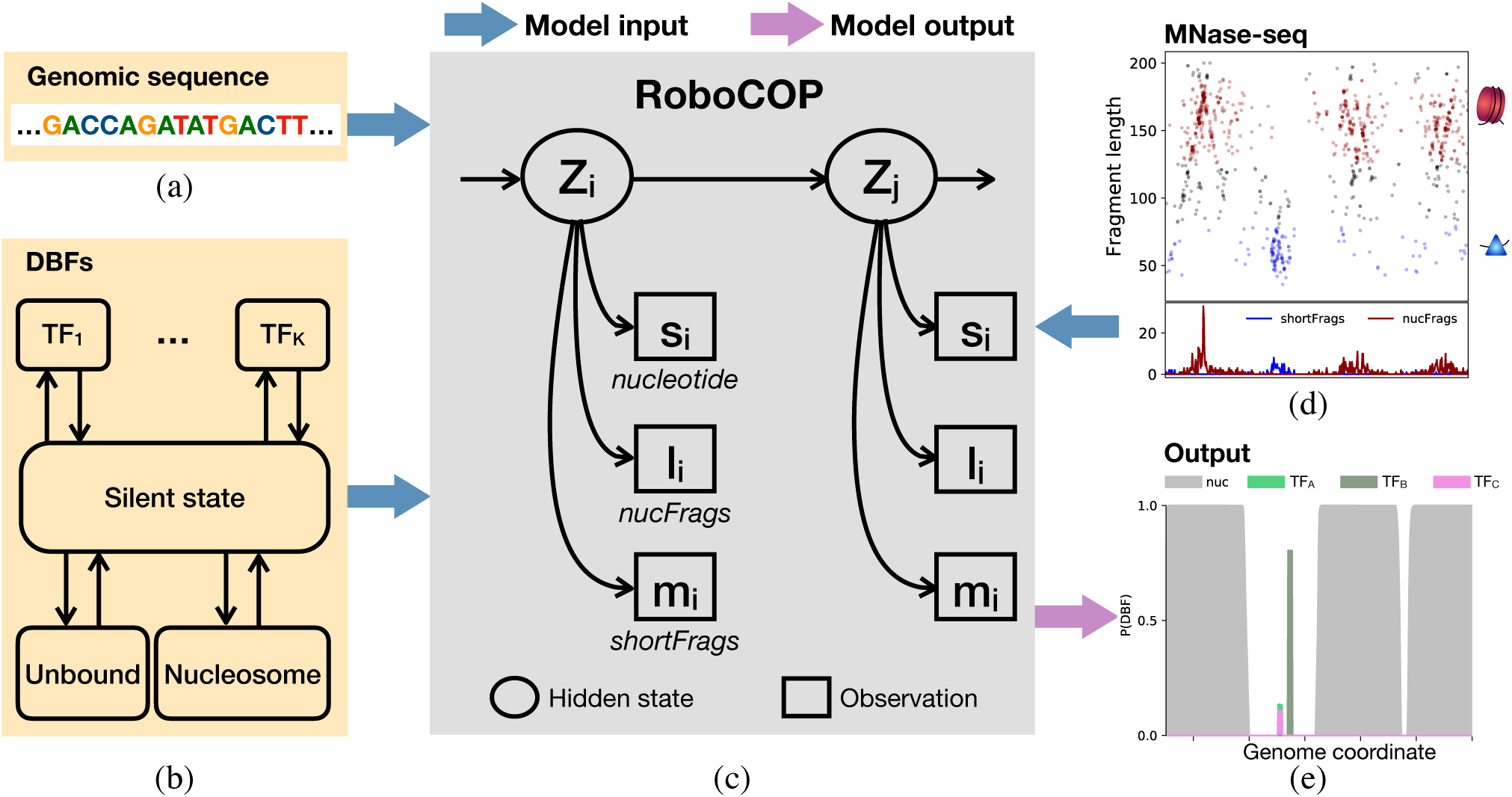
RoboCOP takes various inputs (blue arrows) and produces as output (pink arrow) a chromatin occupancy profile providing quantitative estimates of occupancy for the specified collection of DBFs. The underlying genomic sequence (a) and the collection of DBFs and their sequence specificity models (b) are provided as input to the RoboCOP model (c), along with the `nucFrags` and `shortFrags` signals that result from aggregation of MNase-seq fragment midpoint counts (d). (b) The state transition matrix for the HMM is simplified by the inclusion of a central, non-emitting silent state; from this state, the model can transition to any DBF, after which it necessarily transitions back to the central silent state, thereby removing dependencies among the DBFs. (c) RoboCOP is a multivariate HMM where the hidden state *z*_*i*_ at genomic position *i* emits a nucleotide (*s*_*i*_), a `nucFrags` count (*l*_*i*_), and a `shortFrags` count (*m*_*i*_). (e) RoboCOP performs posterior decoding and yields the probability of each DBF at every position in the genome. The score on the *y*-axis is the probability of that location being bound by a given DBF.

Each genome coordinate is represented by one hidden state in the HMM. An unbound DNA nucleotide is length one, so its model ***π***_0_ has just a single hidden state. The other DBFs (nucleosomes and TFs) have binding sites of greater length and are thus modeled using collections of multiple hidden states. For TF *k* with a binding site of length *L*_*k*_, the HMM either transitions through *L*_*k*_ hidden states of its binding motif or *L*_*k*_ hidden states of the reverse complement of its binding motif. An additional non-emitting state is added as the first hidden state of the TF model ***π***_*k*_, allowing the HMM to transition through the forward or reverse complement of the motif with equal probability (Fig. S2a). The complete TF model ***π***_*k*_ therefore has a total of 2*L*_*k*_ + 1 hidden states. Once the HMM enters the hidden states for either the forward or reverse motif, it transitions through the sequence of hidden states with probability one between consecutive hidden states. On reaching the final hidden state of either motif, the HMM transitions back to the central silent state with probability one. Likewise, once the HMM enters the nucleosome model ***π***_*K*+1_, it transitions through a sequence of hidden states corresponding to 147 nucleotides, after which it transitions back to the central silent state (Fig. S2b). The nucleosome model differs from the TF models in that the latter are modeled with simple PWM motifs, while the former is implemented using a dinucleotide sequence specificity model.

Suppose the sequence of hidden states for the entire genome of length *G* is denoted as *z*_1_, …, *z*_*G*_. Then the transition probabilities satisfy the following:

- *P* (*z*_*g*+1_ = *π*_*k,l*+1_|*z*_*g*_ = *π*_*k,l*_) = 1 whenever *l < L*_*k*_. Within a DBF, the HMM only transitions to that DBF’s next state and not any other state, until it reaches the end of the DBF.
- 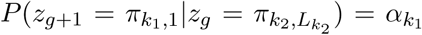 for all *k*_1_ and *k*_2_. The transition probability to the first state of a DBF is a constant, independent of which DBF the HMM visited previously.
- *P* (*z*_*g*+1_|*z*_*g*_) = 0 for all other cases.

The HMM always starts in the central silent state with probability one; this guarantees that it cannot start in the middle of a DBF.

### 2.3 RoboCOP emission probabilities

The HMM employed by RoboCOP is multivariate, meaning that each hidden state is responsible for emitting multiple observables per position in the genome (Fig. 2c). In our case, these observables are modeled as independent, conditioned on the hidden state, but adding dependence would be straightforward. In this paper, we analyze paired-end MNase-seq data, so for a genome of length *G*, the sequences of observables being explained by the model are: (i) nucleotide sequence {*s*_1_, …, *s*_*G*_}, (ii) midpoint counts of MNase-seq `nucFrags` {*l*_1_, …, *l*_*G*_}, and (iii) midpoint counts of MNase-seq `shortFrags` {*m*_1_, …, *m*_*G*_}. For any position *g* in the genome, the hidden state *z*_*g*_ is thus responsible for emitting a nucleotide *s*_*g*_, a number *l*_*g*_ of midpoints of `nucFrags`, and a number *m*_*g*_ of midpoints of `shortFrags` (Fig. 2c). Since these three observations are independent of one another given the hidden state *z*_*g*_, each hidden state has an emission model for each of the three observables, and the joint probability of the multivariate emission is the product of the emission probabilities of the three observables.

For the TF models ***π***_1_, …, ***π***_*K*_, emission probabilities for nucleotide sequences are represented using PWMs. For each of our 150 TFs, we use the PWM of its primary motif reported in (23) (except for Rap1, where we use the more detailed motifs in (5)). For the nucleosome model ***π***_*K*+1_, the emission probability for a nucleotide sequence of length 147 can be represented using a position-specific dinucleotide model (24). To represent this dinucleotide model, the number of hidden states in ***π***_*K*+1_ is roughly 4×147. We use the same dinucleotide model that was used earlier in COMPETE (14).

As described earlier, the two-dimensional MNase-seq data are transformed into two onedimensional signals (Fig. 2d); the midpoint counts of `nucFrags` primarily influence the learned nucleosome positions and the midpoint counts of `shortFrags` primarily influence the learned TF binding sites. In both cases, a negative binomial (NB) distribution is used to model the emission probabilities. We use two sets of NB distributions to model the midpoint counts of `nucFrags`. One distribution, *NB*(***µ***_*nuc*_, *ϕ*_*nuc*_), explains the counts of `nucFrags` at the nucleosome positions and another distribution, *NB*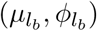, explains the counts of `nucFrags` elsewhere in the genome. Since the midpoint counts of `nucFrags` within a nucleosome are not uniform (Fig. 1b), we model each of the 147 positions separately. To obtain ***µ***_*nuc*_ and *ϕ*_*nuc*_, we collect the midpoint counts of `nucFrags` in a window of size 147 centered on the annotated nucleosome dyads of the top 2000 well-positioned nucleosomes (17) and estimate 147 NB distributions using maximum likelihood estimation (MLE). The 147 estimated values of *µ* are denoted as ***µ***_*nuc*_. The mean of the 147 estimated values of *ϕ* is denoted as *ϕ*_*nuc*_ (shared across all 147 positions). Quantile-quantile plots show the resulting NB distributions to be a good fit (Fig. S3). As for *NB*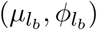, we use MLE to estimate its parameters from the midpoint counts of `nucFrags` within the linker regions on both sides of the same set of 2000 nucleosomes. For this purpose, we considered linkers to be 15 bases long (25).

Similarly, we model the midpoint counts of `shortFrags` using two distributions where one of them, *NB*(*µ*_*T F*_, *ϕ*_*T F*_), explains the counts of `shortFrags` within TF binding sites, while the other, *NB*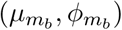, explains counts elsewhere. To estimate the parameters of *NB*(*µ*_*T F*_, *ϕ*_*T F*_), we collect the midpoint counts of `shortFrags` within annotated Abf1 and Reb1 binding sites (18) and fit the NB distribution using MLE. A quantile-quantile plot again shows the NB distribution provides a good fit (Fig. S4). We chose Abf1 and Reb1 for fitting the distribution because these TFs have many binding sites in the genome and the binding sites are often less noisy. For parameterizing *NB*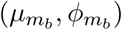, we collect the midpoint counts of `shortFrags` within the same linker regions used earlier and estimate the NB distribution using MLE.

### 2.4 RoboCOP transition probability updates

Within each single DBF model, the transition probabilities between hidden states can only be zero or one (except for the two transition probabilities from each TF model’s first, nonemitting state to the first state of either its forward or reverse motif; these are fixed at 0.5.) Consequently, the only transition probabilities we need to learn are {*α*_0_, …, *α*_*K*+1_}, those from the central silent state to the first state of each DBF (Fig. S1). Our approach is to initialize these to sensible values, and then optimize them using Baum-Welch, which is guaranteed to converge to a local maximum of the model’s likelihood.

To initialize the transition probabilities {*α*_0_, …, *α*_*K*+1_}, we first assign a non-negative concentration or ‘weight’ to each DBF. Let the weight for DBF *i* be denoted *w*_*i*_. Following previous work (14; 15), we assign weight *w*_0_ = 1 to the ‘empty’ DBF (representing an unbound DNA nucleotide) and *w*_*K*+1_ = 35 to the nucleosome. To each TF *k* ∈ {1, …, *K*}, we assign a weight *w*_*k*_ which is that TF’s dissociation constant *K*_*D*_ (or alternatively, a multiple thereof: 8*K*_*D*_, 16*K*_*D*_, 32*K*_*D*_, or 64*K*_*D*_).

To convert these weights into transition probabilities, we need to compensate for the fact that each DBF *k* has a different length *L*_*k*_, from as little as one, for an unbound nucleotide, to 147, for a nucleosome. A little algebra, and the fact that *w*_0_ = 1, allows us to write the following relationship to account for the difference in lengths:

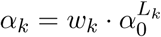

Since {*α*_0_, …, *α*_*K*+1_} are a set of probabilities, it must also be the case that they sum to 1:

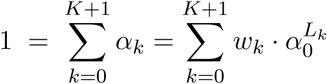

Finally, because we know all the values of *w*_*k*_ and *L*_*k*_, we are left with an expression in just one unknown, *α*_0_. We can easily solve for *α*_0_, and then use it and the relationship above to compute the transition probabilities of all the other DBFs.

After initializing the transition probabilities as described above, we iteratively update them using Baum-Welch until convergence to a local optimum of the likelihood. To update *α*_*k*_, we compute:

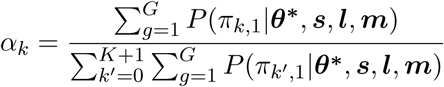

Here, ***θ***^*^ represents all the model parameters. We find the likelihood converges within ten iterations (Fig. S5) and the optimized transition probabilities for each DBF almost always converge to the same final values regardless of how we initialize the weights (Fig. S6). We find convergence is faster for most DBFs when we initialize TF weights to *K*_*D*_ rather than multiples thereof (Fig. S6).

We find that transition probabilities for a few TFs with AT-rich motifs like Azf1 and Smp1 can grow quite large, resulting in a large number of binding sites in the genome, most of which are potential false positives. To curb the number of binding site predictions for such TFs, we apply a threshold on TF transition probabilities. The threshold *δ* is chosen to be two standard deviations above the mean of the initial transition probabilities of all the TFs (Fig. S7). Therefore, after the Baum-Welch step in every iteration, an additional modified Baum-Welch step is computed as follows:

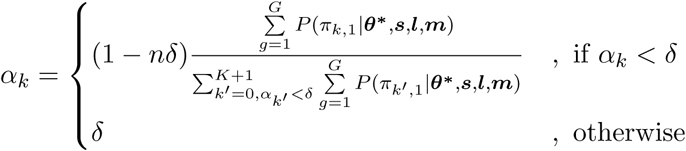

where *n* is the number of TFs that have a transition probability more than *δ*. So, for all the TFs whose transition probabilities would be more than *δ*, they are instead set to *δ*, and the remaining DBFs (including the nucleosome and unbound state) have a regular Baum-Welch update of their transition probabilities. We find that this approach reduces the number of false positives (Fig. S8). An alternative mechanism might be to use an informed prior, in situations where prior information is available.

Note that when we compare RoboCOP and COMPETE, we run COMPETE with the exact same model parameters as RoboCOP in order to isolate the differences in the output profiles that arise from the inclusion of chromatin accessibility data as input to RoboCOP. Because model parameters include DBF transition probabilities, and because RoboCOP has access to chromatin accessibility data when it estimates these with Baum-Welch, this potentially gives COMPETE a slight advantage it would not normally have.

### 2.5 Implementation details for posterior decoding

RoboCOP employs posterior decoding to infer probabilistic occupancy profiles of protein-DNA binding. The motivation behind posterior decoding is that it represents the thermodynamic ensemble of potential binding configurations; the resulting probability distribution sheds light on the many different ways proteins may be bound to the genome across a cell population (applying Viterbi decoding would not provide a probabilistic landscape, but only a single, most likely chromatin configuration). The resulting posterior probability of each DBF at each position in the genome provides a probabilistic profile of DBF occupancy at base-pair resolution (Fig. 2e).

As a multivariate HMM, RoboCOP has a time complexity of *O*(*GN* ^2^) and a space complexity of *O*(*GN*) (for a genome of length *G* and where *N* denotes the total number of hidden states). The high complexity makes it difficult to decode the entire genome at once. To reduce the computational complexity of RoboCOP, we perform posterior decoding separately on blocks of the genome of length 5000, with an overlap of 1000 bases, and stitch results together. This ensures that the model has a sufficiently long sequence to learn an accurate chromatin landscape, but not so long that we run out of memory. In addition, we use only the longest chromosome (chrIV in yeast) to train DBF transition probabilities with Baum-Welch, and then undertake posterior decoding genome-wide.

### 2.6 Validation of TF and nucleosome predictions

We use posterior probabilities of TF occupancy from RoboCOP and COMPETE output to identify binding sites, calling all sites whose starting probability is at least 0.1. The starting probability of a motif is computed by adding the starting probability of the forward and reverse complement of the motif for every position in the genome. In the case of Rap1 which has multiple PWMs, the maximum starting probability among the PWMs is chosen at every position. For validation, a site is considered a true positive (TP) if it overlaps with an annotated binding site for that TF, and a false positive (FP) otherwise. If an annotated TF binding site does not overlap any of our predictions, it is a false negative (FN).

We call nucleosomes from RoboCOP and COMPETE outputs using a greedy algorithm, as described previously (26). Briefly, nucleosome dyads with decreasing probability are iteratively selected. A window of size 101 around the selected dyad is removed from future rounds of dyad selection (this window size is chosen to allow mild overlap between adjacent nucleosome locations). The annotations from Brogaard and colleagues (17) contain 67548 nucleosomes. We select the same number of nucleosomes from the outputs of RoboCOP and COMPETE, respectively. For validation, a nucleosome position is considered a true positive (TP) if the distance between the predicted and annotated dyad is less than 50 bases.

### 2.7 FIMO-MNase

To calibrate the accuracy of the TF binding site predictions of RoboCOP and COMPETE, we developed a baseline by running FIMO (27) on non-occluded peaks of the `shortFrags` signal as follows. We first smooth MNase-seq midpoint counts of `shortFrags` using a window of size 21. Then, we call a peak if its height is greater than 2 and it is at least 25 bases away from any other peak. We call nucleosomal peaks using the midpoint counts of `nucFrags` when their height is greater than 1 and they are at least 100 bases apart. Finally, to prevent nucleosomal peaks occluding peaks of `shortFrags`, we remove peaks of `shortFrags` that are within 60 bases of peaks of `nucFrags`. After these steps, we detect 4137 non-occluded peaks of `shortFrags` genome-wide. Within 50-bp windows centered on these peaks, we use FIMO (27) to scan for matches to any of our PWMs, with a p-value cutoff of 10^−4^.

### 2.8 Open data and software access

MNase-seq and RNA-seq data from yeast cells before and after 60 minutes of cadmium treatment are already available from the Duke Digital Repository at https://doi.org/10.7924/r4hx1b43s and these data will also be uploaded to GEO prior to publication. The repository contains a snapshot of our RoboCOP code, but the latest version can always be downloaded from https://github.com/HarteminkLab/RoboCOP. We use the sacCer2 (June 2008) version of the yeast genome for all our analyses.

## 3 RESULTS

### 3.1 RoboCOP computes probabilistic chromatin occupancy profiles

We use RoboCOP to predict the nucleosome positions and binding sites of 150 different TFs across the *Saccharomyces cerevisiae* genome. Even though we include 150 different TFs in our model (listed in Table S1), this does not exhaust what binds the genome: We are missing replication factors and general transcription factors, as well as sequence-specific TFs for which we have no binding preference information. To address this, we add a 10-bp DBF labeled ‘unknown’ that we use to capture any extra `shortFrags` signal not captured by our 150 known TFs (this also has the salutary effect of reducing false positive predictions for the known TFs; see Fig. S8 for a comparison).

Beyond the genome sequence and the collection of DBFs and their binding preferences, RoboCOP takes as input the `nucFrags` and `shortFrags` signals derived from paired-end MNase-seq data. Fig. 3 shows the input MNase-seq data and the resulting RoboCOP output for a representative segment of the genome. The nucleosome predictions in RoboCOP’s output (Fig. 3c) line up well with the nucleosomal fragments in the data (Fig. 3a,b). In addition, RoboCOP predicts one Abf1 and one Reb1 binding site, which align with the short fragments in the data and match annotated binding sites in this locus (18).

**Figure 3.**
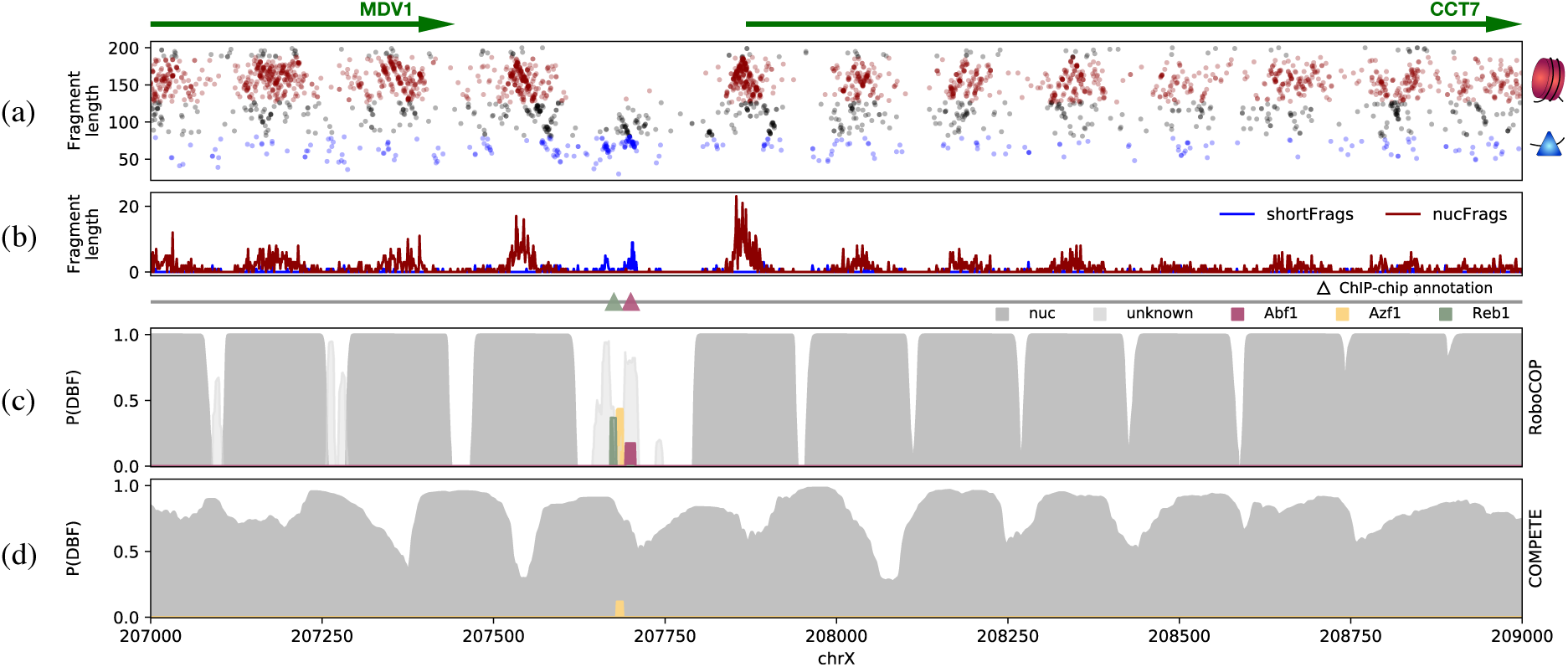
Representative chromatin occupancy profile produced by RoboCOP, in comparison with that of COMPETE, an existing method. (a) Two-dimensional plot of MNase-seq fragments from positions 207,000 to 209,000 of yeast chromosome X, with `nucFrags` in red and `shortFrags` in blue. Gene annotations depicted with arrows at the top. (b) The `nucFrags` and `shortFrags` signals that result from aggregation of MNase-seq fragment midpoint counts in the region. (c) RoboCOP and (d) COMPETE outputs for the region, with known TF binding sites indicated with triangles above. Because RoboCOP makes use of MNase-seq data in generating its chromatin occupancy profile, it, unlike COMPETE, positions nucleosomes more precisely and successfully identifies not only the nucleosome-depleted region, but also the known Abf1 and Reb1 binding sites therein (18).

### 3.2 RoboCOP’s use of chromatin accessibility data improves chromatin occupancy profiles

Our group’s earlier work, COMPETE (14) uses only nucleotide sequence as input to an HMM in order to compute a probabilistic occupancy landscape of DBFs across a genome. COMPETE’s output is theoretical in that it does not incorporate experimental data in learning the binding landscape of the genome. Perhaps unsurprisingly, the nucleosome positions learned by COMPETE (Fig. 3d) do not line up well with the nucleosomal signal apparent in the MNase-seq data (Figs. 3a,b). The nucleosome predictions of COMPETE (Fig. 3d) are more diffuse, which is understandable because it relies entirely on sequence information, and nucleosomes have only weak and periodic sequence specificity (1). Because of a lack of chromatin accessibility data, COMPETE fails to identify the clear nucleosome-depleted region in this locus (and does so all throughout the genome, as seen in Figs. 4a,b), as a result of which it fails to recognize the Abf1 and Reb1 binding sites known to reside in the locus in (18). In contrast, RoboCOP utilizes the chromatin accessibility data to accurately learn the nucleosome positions and the annotated Abf1 and Reb1 binding sites (Fig. 3c).

**Figure 4.**
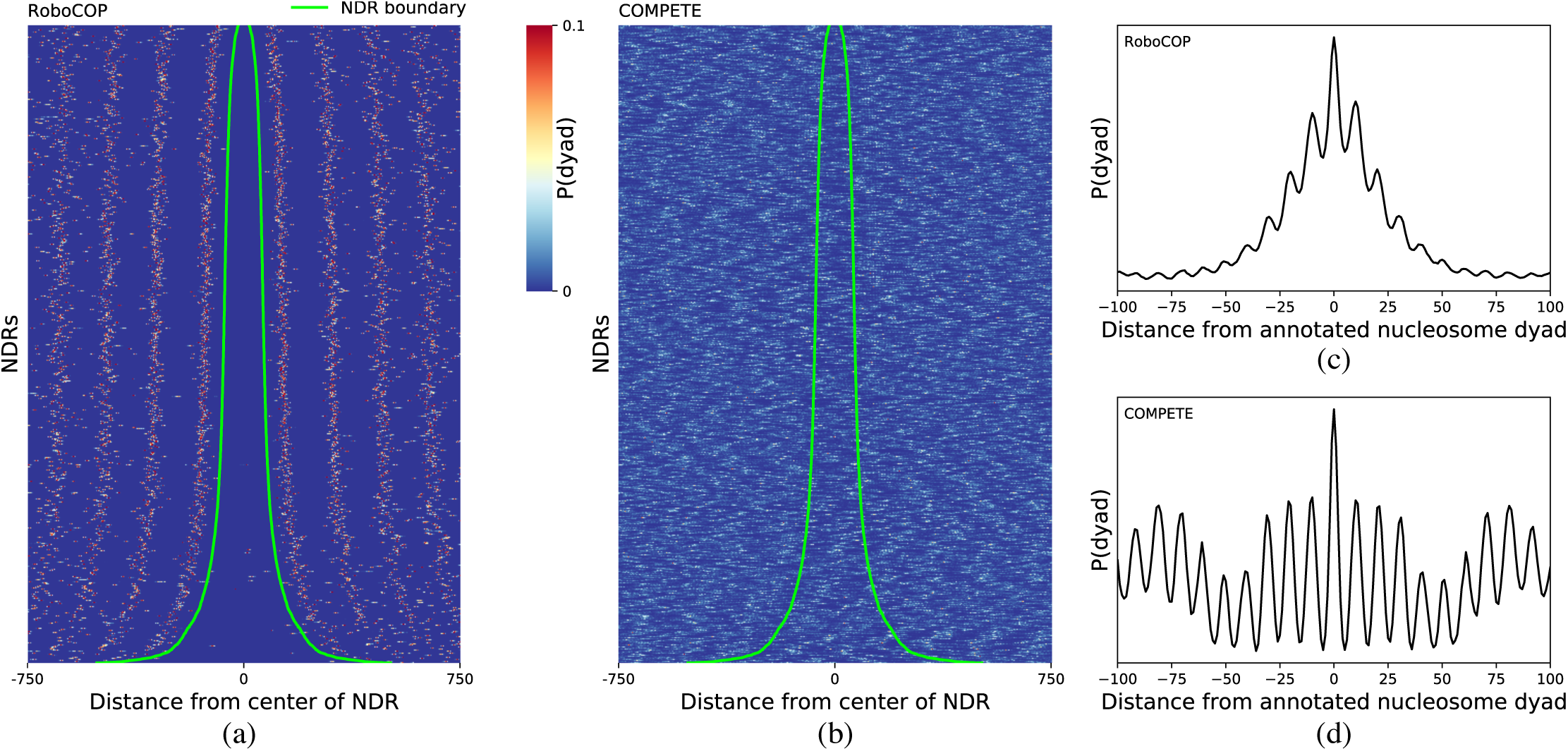
RoboCOP positions nucleosomes with precision and accuracy, including avoiding their placement within nucleosome-depleted regions (NDRs). (a,b) Heatmaps depict the predicted probability of a nucleosome dyad, P(dyad), at each position around experimentally determined NDRs genome-wide (25), as computed by (a) RoboCOP and (b) compete. Each row is a distinct NDR, sorted by NDR size. Lime green lines depict the experimentally determined NDR boundaries. Note that P(dyad) computed by RoboCOP is appropriately almost always zero within NDRs, unlike COMPETE, and the signal is well-phased in both directions. (c,d) Curves depict aggregate values of P(dyad) across all annotated nucleosome dyads genome-wide (17), as computed by (c) RoboCOP and (d) COMPETE. Both aggregate signals have an expected ∼10 bp periodicity that arises from the periodic nature of the weak sequence specificity of nucleosomes. Note that P(dyad) computed by RoboCOP appropriately peaks at annotated dyads and falls off rapidly in both directions, indicating that learned positions are both more precise and more accurate than those of COMPETE.

### 3.3 Predicted nucleosome positions

Nucleosomes have weak sequence specificity and can adopt alternative nearby positions along the genome (25; 28). It is therefore likely that the nucleosome positions reported by one method do not exactly match those reported by another. However, since RoboCOP generates genome-wide probabilistic scores of nucleosome occupancy, we can plot the probability of a nucleosome dyad, P(dyad), around annotated nucleosome locations (17). We find that the RoboCOP dyad score peaks precisely at the annotated dyads (Fig. 4d), and decreases almost symmetrically in either direction. In contrast, COMPETE does not provide accurate location predictions (Fig. 4c); the oscillatory nature of the score reported by COMPETE reflects the periodic dinucleotide sequence specificity model for nucleosomes, and does not correspond well with actual nucleosome locations. When evaluated genome-wide using an F1-score to balance the trade-off between precision and recall (Fig. 5), the nucleosome positions called by RoboCOP are far more similar to the nucleosome annotations of Brogaard and colleagues (17) than are the ones called by COMPETE, which are only slightly better than random (Table S3).

**Figure 5.**
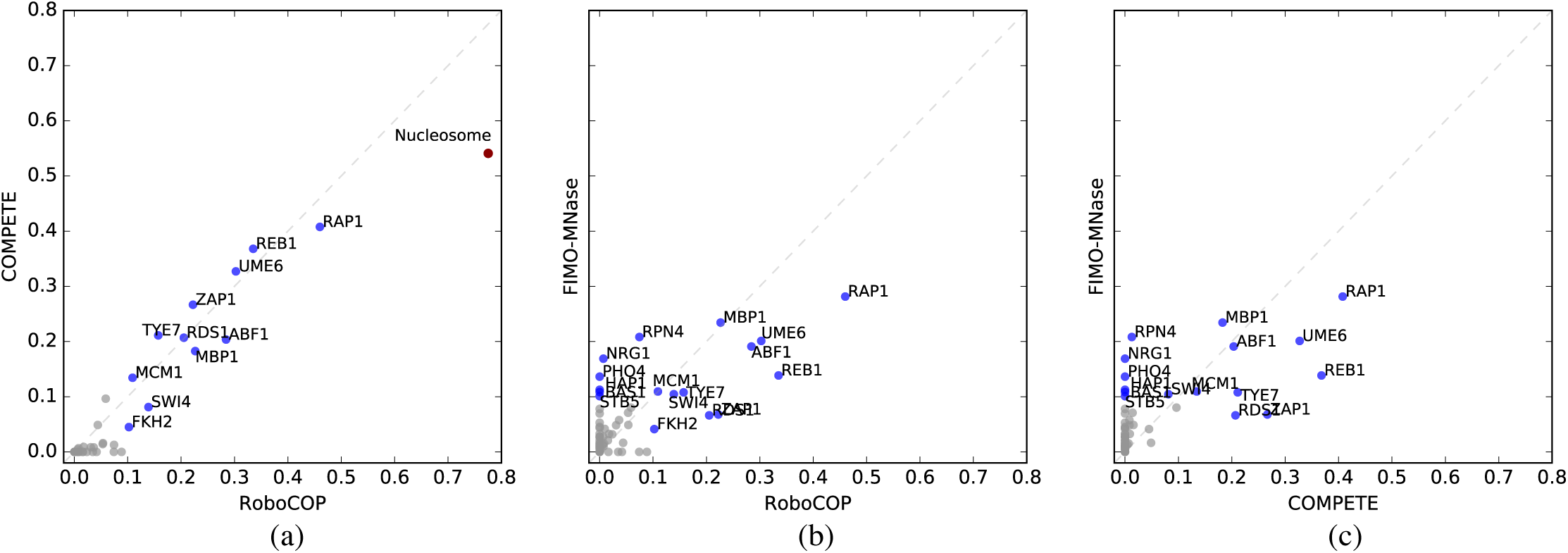
(a) In comparing the F1-scores of the genome-wide predictions made by RoboCOP and COMPETE, RoboCOP does mildly better on TF binding site predictions and markedly better on nucleosome predictions. TFs with F1-score less than 0.1 in both methods are colored gray. As a baseline, we also compare the F1-scores of the genome-wide TF binding site predictions of (b) RoboCOP and FIMO-MNase, and (c) COMPETE and FIMO-MNase.

### 3.4 Predicted TF binding sites

MNase-seq is primarily used to study nucleosome positions; at present, no methods exist to predict TF binding sites from MNase-seq. It is also challenging to extract TF binding sites from the noisy `shortFrags` signal that results from MNase digestion. TFs can sometimes be bound for an extremely short span of time (29), allowing the entire region to be digested by MNase and leaving behind no `shortFrags` signal. Nevertheless, MNase-seq data has been reported to provide evidence of binding for at least some TFs and DNA replication initiation factors (12; 30; 13), so we explored how well RoboCOP is able to identify TF binding sites.

Although RoboCOP predicts the genome-wide occupancy of a set of 150 TFs, we can only validate the binding sites of 81 of them, given available ChIP-chip (18), ChIP-exo (5), and ORGANIC (19) datasets (Table S1). Making things more complicated, available yeast ChIP-chip data assay binding at the genomic resolution of whole intergenic regions, with computational algorithms being used to refine those into specific binding sites, making the ChIP-chip dataset somewhat less reliable for validation purposes. Compounding the problem, data for many of the TFs were generated under multiple conditions (3) (Table S1) and these conditions are not specified as part of the annotations.

With those caveats in place, we compare TF binding site predictions made by RoboCOP to predictions made by COMPETE and observe mild but consistent improvement in F1-scores with RoboCOP (Fig. 5a). As a baseline for these two methods, we compare their results to an approach we call FIMO-MNase, in which we run FIMO (27) around the peaks of midpoint counts of MNase `shortFrags`. We find RoboCOP and COMPETE generally perform better than FIMO-MNase, although they both have difficulty with a few factors (Figs. 5b,c). We have the most precise binding site annotation datasets for Abf1, Reb1, and Rap1, and for these TFs, both COMPETE and RoboCOP make markedly better predictions than FIMO-MNase. Overall, the highest F1-score is for Rap1 binding site predictions made by RoboCOP.

### 3.5 RoboCOP reveals chromatin dynamics under cadmium stress

One of the most powerful uses of RoboCOP is that it can elucidate the dynamics of chromatin occupancy, generating profiles under changing environmental conditions. As an example, we explore the occupancy profiles of yeast cells before and after being subjected to cadmium stress for 60 minutes. We run RoboCOP separately on two MNase-seq datasets: one for a cell population before treatment and another 60 minutes after treatment with 1mM of CdCl_2_. Cadmium is toxic to the cells and activates stress response pathways. Stress response genes are heavily transcribed under cadmium treatment, while ribosomal genes are repressed (31). We use RNA-seq to identify the 100 genes most up-regulated (‘upmost 100’, for short) and the 100 genes most down-regulated (‘downmost 100’). As a control, we choose the 100 genes with the least change in transcription under treatment (‘constant 100’) (see Table S2 for the three gene lists). Separately for each group of genes, we plot the composite RoboCOP-predicted nucleosome dyad probability in a 1000-bp window centered on established +1 nucleosome annotations (25). Prior to cadmium treatment, the composite +1 nucleosome peaks for all three groups align closely with the annotations (filled curves in Figs. 6 a,b,c). Upon treatment with cadmium, the +1 nucleosomes of the upmost 100 genes shift downstream, expanding the NDR (solid curve in Fig. 6a). Owing to high variability in the new positions of the +1 nucleosomes of the upmost 100 genes, the composite +1 nucleosome peak for these genes becomes shorter and broader. Furthermore, the position of the −1 nucleosome also becomes more uncertain with the expansion of the NDR. In contrast, the +1 nucleosomes of the downmost 100 genes shift upstream, closing in on the NDR (solid curve in Fig. 6c). Interestingly, the shift is precise, resulting in the composite +1 nucleosome peak remaining narrow and sharp. Unlike the upmost100 genes, we do not see changes in the position of the −1 nucleosomes of the downmost 100 genes. As expected from a control, we observe no changes in the position of the +1 nucleosome for the constant 100 genes (Fig. 6b).

**Figure 6.**
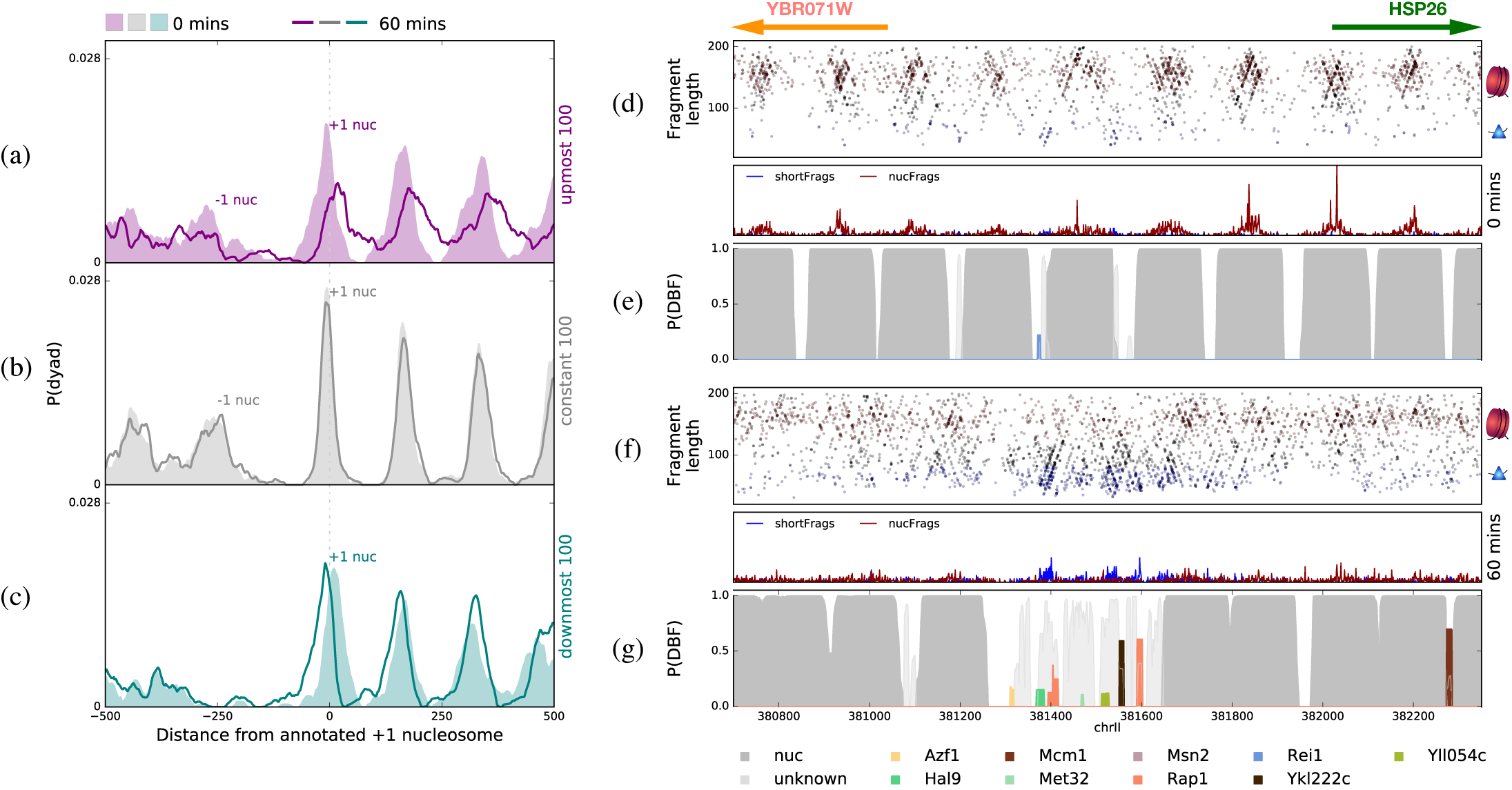
(a–c) Aggregate nucleosome dyad probability, as computed by RoboCOP, around annotated +1 nucleosomes (25) of (a) the 100 most up-regulated genes (purple), (b) the 100 genes least changed in transcription (gray), and (c) the 100 most down-regulated genes (teal), before and 60 minutes after treating cells with cadmium. After treatment, we see the +1 nucleosome closing in on the promoters of the most down-regulated genes (teal) but opening up the promoters of the most up-regulated genes (purple). (d) Two-dimensional plot of MNase-seq fragments near the HSP26 promoter (positions 380,700 to 382,350 of yeast chromosome II are shown) before treatment with cadmium (`nucFrags` in red; `shortFrags` in blue), along with the `nucFrags` and `shortFrags` signals that result from aggregating those midpoint counts. Gene annotations depicted with arrows at the top (Watson strand in green; Crick strand in orange). (e) RoboCOP-predicted occupancy profile of this region before treatment with cadmium. (f,g) The same as (d,e), respectively, but 60 minutes after cadmium treatment. HSP26 transcription is highly up-regulated under cadmium stress, and we observe here that its promoter exhibits marked TF binding after treatment, most prominently by Rap1, known to bind this promoter during stress response. Nucleosome positions also shift notably.

We can also use RoboCOP to study detailed changes in the chromatin landscape under cadmium stress within a specific locus, for example that of HSP26, a key stress response gene in the upmost 100 genes. In Figs. 6d-g, we notice the HSP26 promoter opening up under stress, with shifts in nucleosomes leading to more TF binding in the promoter. From the `shortFrags` midpoint counts, RoboCOP identifies multiple potential TF binding sites, most prominently for Rap1, which has already been shown to re-localize to the promoter region of HSP26 during general stress response (32).

In comparison, COMPETE fails to capture the dynamics of chromatin occupancy under cadmium stress because it does not incorporate chromatin accessibility information into its model. We ran COMPETE with the RoboCOP-trained DBF weights for the two time points of cadmium treatment and found that COMPETE generates binding landscapes for the two time points that are nearly identical (Fig. S9). This is a key difference between RoboCOP and COMPETE: Being able to incorporate experimental chromatin accessibility data allows RoboCOP to provide a more accurate binding profile for cell populations undergoing dramatic chromatin changes.

The preceding analysis highlights the broad utility of RoboCOP. Because RoboCOP models DBFs competing to bind the genome, it produces a probabilistic prediction of the occupancy level of each DBF at single-nucleotide resolution. Moreover, as the chromatin architecture changes under different environmental conditions, RoboCOP is able to elucidate the dynamics of chromatin occupancy. The cadmium treatment experiment shows that the predictions made by RoboCOP can be used both to study overall changes for groups of genes (Figs. 6a-c), as well as to focus on specific genomic loci in order to understand their detailed chromatin dynamics (Fig. 6d-g).

## 4 DISCUSSION

RoboCOP is a new computational method that utilizes a multivariate HMM to generate a probabilistic occupancy profile of the genome by integrating chromatin accessibility data with nucleotide sequence. We chose to apply the model to the yeast genome because of the availability of high quality MNase-seq data and the small size of the genome, which simplifies computation. Chromatin accessibility data from MNase-seq, DNase-seq, and ATAC-seq are generally noisy, so it is a challenging task to infer precise genome-wide DBF occupancy from the data, particularly for TFs. While alternative approaches using peak or footprint identification followed by TF-labeling with FIMO (27) can offer some insight into protein-DNA binding, we observe that RoboCOP performs notably better, presumably because it considers all DBFs together within a single joint model that explicitly accounts for the thermodynamic competition among DBFs, including nucleosomes.

RoboCOP improves upon COMPETE in a number of ways: It increases the accuracy of TF binding site predictions, it markedly increases the accuracy of nucleosome positioning predictions, and it uses experimental data to learn DBF transition probabilities in a principled way. When these same transition probabilities are provided to COMPETE, its TF binding site predictions are fairly similar to RoboCOP’s because of the generally high sequence specificity of TFs, but its nucleosome positioning predictions are much worse because of the weak sequence specificity of nucleosomes. In future work, it might be possible to further improve the TF binding site predictions and the estimated transition probabilities through the incorporation of prior information.

In closing, we note that RoboCOP can be used to study the chromatin architecture of the genome under varying conditions, a task to which COMPETE is unsuited. Because RoboCOP uses chromatin accessibility data in its model of DBFs competing to bind to the genome, it is able to reveal dynamic levels of occupancy for each DBF at each location throughout the genome as environmental conditions change. Importantly, since gene expression also varies in response to changing environmental conditions, we believe RoboCOP will help elucidate how the dynamics of chromatin occupancy and the dynamics of gene expression interrelate.

## ACKNOWLEDGMENTS

The authors would like to thank Heather MacAlpine and Vinay Tripuraneni for generating the MNase-seq data, and Greg Crawford, Raluca Gordân, Ed Iversen, Trung Tran, Yulong Li, and Albert Xue for helpful comments and feedback during the development of RoboCOP. This work was supported by the following grants from the National Institute of General Medical Sciences: R35 GM127062 (D.M.M.) and R01 GM118551 (A.J.H.).

