## Supplementary Material for "RoboCOP: Jointly computing chromatin occupancy profiles for numerous factors from chromatin accessibility data"

May 12, 2020

| | | | | | |
|---|---|---|---|---|---|
| Abf1 | Abf2 | Ace2 | Adr1 | Aft1 | Aft2 |
| Aro80 | Asg1 | Azf1 | Bas1 | Cad1 | Cat8 |
| Cbf1 | Cep3 | Cha4 | Cin5 | Crz1 | Cst6 |
| Cup9 | Dal80 | Dal82 | Ecm22 | Ecm23 | Fhl1 |
| Fkh1 | Fkh2 | Fzf1 | Gal4 | Gat1 | Gat3 |
| Gat4 | Gcn4 | Gcr1 | Gis1 | Gln3 | Gsm1 |
| Gzf3 | Hac1 | Hal9 | Hap1 | Hcm1 | Hmlalpha2 |
| Hmra2 | Hsf1 | Leu3 | Lys14 | Matalpha2 | Mbp1 |
| Mcm1 | Met31 | Met32 | Mga1 | Mig1 | Mig2 |
| Mig3 | Mot3 | Msn1 | Msn2 | Msn4 | Ndt80 |
| Nhp10 | Nhp6a | Nhp6b | Nrg1 | Nrg2 | Oaf1 |
| Pbf1 | Pbf2 | Pdr1 | Pdr3 | Phd1 | Pdr8 |
| Pho2 | Pho4 | Put3 | Rap1 | Rdr1 | Rds1 |
| Rds2 | Reb1 | Rei1 | Rfx1 | Rgm1 | Rgt1 |
| Rim101 | Rox1 | Rph1 | Rpn4 | Rsc3 | Rsc30 |
| Rtg3 | Sfl1 | Sfp1 | Sig1 | Sip4 | Skn7 |
| Sko1 | Smp1 | Sok2 | Spt15 | Srd1 | Stb3 |
| Stb4 | Stb5 | Ste12 | Stp1 | Stp2 | Stp3 |
| Stp4 | Sum1 | Sut1 | Sut2 | Swi4 | Swi5 |
| Tbf1 | Tbs1 | Tea1 | Tec1 | Tos8 | Tye7 |
| Uga3 | Ume6 | Upc2 | Usv1 | Vhr1 | Xbp1 |
| Yap1 | Yap3 | Yap6 | YBR033W | YBR239C | YDR520C |
| YER064C | YER130C | YER184C | YGR067C | YKL222C | YLL054C |
| YLR278C | YML081W | YNR063W | Yox1 | YPR013C | YPR015C |
| YPR022C | YPR196W | Yrm1 | Yrr1 | Zap1 | Zms1 |

Table S1: List of 150 TFs used by RoboCOP. TFs ones colored in gray do not have binding sites reported in (1). TFs colored in red have alternative validation sets from ChIP-exo (2) and ORGANIC (3) protocols. TFs colored in blue are known to be expressed at increased levels under specific experimental conditions, so may not be expressed in our own data (4).

| upmost 100 | downmost 100 | constant 100 |
|---|---|---|
| HSP12 (6.30) | RPL9A (−4.59) | AIF1 (0.00) |
| MET3 (5.87) | RPS22A (−4.51) | YMR272W-B (0.00) |
| PDC6 (5.69) | SSB1 (−4.43) | YBR013C (0.00) |
| YDL124W (5.59) | RPS31 (−4.42) | YNR070W (0.00) |
| HSP26 (5.41) | RPS3 (−4.36) | COS12 (0.00) |
| MET14 (5.41) | ASC1 (−4.34) | UBS1 (0.00) |
| DDR48 (5.15) | RPL3 (−4.34) | IPL1 (0.00) |
| AHP1 (4.99) | YEF3 (−4.31) | SEN54 (0.00) |
| MET10 (4.85) | RPL8B (−4.30) | OSW1 (0.00) |
| STF2 (4.70) | RPS9B (−4.27) | YJL043W (0.00) |
| GRE1 (4.69) | RPL5 (−4.25) | FIG1 (0.00) |
| SPG4 (4.60) | RPS5 (−4.20) | DYN3 (0.00) |
| ECM17 (4.46) | RPP2B (−4.16) | BCK1 (0.00) |
| PRB1 (4.26) | RPS12 (−4.11) | CTF3 (0.00) |
| CYS3 (4.17) | RPL12A (−4.11) | IST2 (0.00) |
| PNC1 (4.09) | RPS2 (−4.01) | VEL1 (0.00) |
| YCT1 (3.99) | RPP2A (−3.97) | YLR406C-A (0.00) |
| BDS1 (3.97) | RPP0 (−3.96) | SMA2 (0.00) |
| PBI2 (3.94) | RPP1A (−3.86) | MEI5 (0.00) |
| ZWF1 (3.92) | RPS15 (−3.82) | MF(ALPHA)1 (0.00) |
| TMA10 (3.92) | RPS0B (−3.81) | MMS1 (0.00) |
| RTC3 (3.90) | RPL28 (−3.80) | ECM12 (0.00) |
| TIS11 (3.84) | RPL38 (−3.79) | HPR1 (0.00) |
| KAR2 (3.79) | RPS14A (−3.75) | PEX25 (0.00) |
| BNA3 (3.78) | ILV5 (−3.75) | NPL6 (0.00) |
| HSP42 (3.71) | RPL16B (−3.72) | YPL009C (0.00) |
| UBI4 (3.68) | RPL22A (−3.71) | ESC8 (0.00) |
| TSA1 (3.67) | RPL2A (−3.70) | RRN5 (0.00) |
| PEP4 (3.67) | RPL1B (−3.68) | YHR214C-E (0.00) |
| LAP4 (3.67) | RPL2B (−3.67) | GET1 (0.00) |
| MET16 (3.60) | RPL32 (−3.64) | SWI3 (0.00) |
| ICY2 (3.59) | SSB2 (−3.64) | GNT1 (0.00) |
| PHO89 (3.58) | RPL1A (−3.64) | YOL019W-A (0.00) |
| YNL208W (3.54) | RPS16B (−3.64) | HIS3 (0.00) |
| MET6 (3.53) | RPL18A (−3.63) | GPI15 (0.00) |
| SEO1 (3.52) | RPL25 (−3.61) | KAR5 (0.00) |
| FIT3 (3.48) | RPS13 (−3.61) | SPT23 (0.00) |
| AGP3 (3.42) | RPL11A (−3.60) | SPT10 (0.00) |
| HSP104 (3.39) | RPL21A (−3.60) | SPR28 (0.00) |
| TDH1 (3.37) | RPL26B (−3.58) | SOG2 (0.00) |
| HBN1 (3.34) | RPL17A (−3.58) | DCG1 (0.00) |
| FIT2 (3.31) | RPL15A (−3.57) | STH1 (0.00) |
| GLK1 (3.30) | RPL30 (−3.57) | THP1 (0.00) |
| MET1 (3.30) | RPS11A (−3.57) | HST4 (0.00) |
| SSA1 (3.29) | RPS8A (−3.54) | EST1 (0.00) |
| AIM17 (3.19) | RPS1B (−3.52) | YKR041W (0.00) |
| YNL134C (3.17) | RPS4A (−3.52) | MRPS5 (0.00) |
| GRE3 (3.11) | RPS26B (−3.51) | OST1 (0.00) |
| PGK1 (3.11) | RPL19B (−3.51) | YDL133W (0.00) |
| HSP82 (3.11) | RPL31A (−3.49) | POP1 (0.00) |
| TSL1 (3.11) | RPS7A (−3.49) | SED5 (0.00) |
| ERO1 (3.09) | SNU13 (−3.47) | MSH4 (0.00) *cont.* |

| upmost 100 | downmost 100 | constant 100 |
|---|---|---|
| PRX1 (3.07) | RPS4B (−3.46) | CCZ1 (0.00) |
| YMR315W (3.04) | RPS20 (−3.44) | SPT8 (0.00) |
| TPS1 (3.03) | RPL42B (−3.44) | MDM1 (0.00) |
| TFS1 (3.00) | RPL27A (−3.42) | HMI1 (0.00) |
| MXR1 (2.99) | RPS18B (−3.42) | VID24 (0.00) |
| PDI1 (2.99) | RPS0A (−3.39) | STB6 (0.00) |
| SAM2 (2.99) | RPL12B (−3.37) | CDC9 (0.00) |
| TRX2 (2.97) | RPS10A (−3.37) | YOR387C (0.00) |
| YBR085C-A (2.97) | RPL13B (−3.36) | SPC110 (0.00) |
| MRP8 (2.97) | RPL6B (−3.36) | STE5 (0.00) |
| FMO1 (2.95) | RPL8A (−3.35) | RTC4 (0.00) |
| MET2 (2.95) | RPL24A (−3.34) | TOP1 (0.00) |
| YOR285W (2.94) | NSR1 (−3.33) | YCR090C (0.00) |
| MUP3 (2.93) | RPS24B (−3.30) | YPR027C (0.00) |
| GSP2 (2.92) | RPS18A (−3.30) | PSR2 (0.00) |
| ALD4 (2.92) | RPS6B (−3.30) | BUB1 (0.01) |
| GRX1 (2.90) | RPL11B (−3.29) | YOR072W-B (0.01) |
| CMK2 (2.88) | RPS6A (−3.28) | LDB16 (0.01) |
| ENO1 (2.88) | RPL6A (−3.26) | YDR042C (0.01) |
| YLR194C (2.86) | RPL20A (−3.23) | ATS1 (0.01) |
| SER33 (2.86) | RPS24A (−3.22) | LOH1 (0.01) |
| ARA1 (2.85) | RPL43A (−3.22) | TAF13 (0.01) |
| GRE2 (2.84) | RPS8B (−3.21) | YER188C-A (0.01) |
| GPD1 (2.83) | RPL33A (−3.21) | MSI1 (0.01) |
| MUP1 (2.82) | NOP1 (−3.20) | YDR194W-A (0.01) |
| HOR7 (2.82) | RPL4A (−3.19) | RUD3 (0.01) |
| DDR2 (2.82) | RPL14A (−3.17) | YHR100C (0.01) |
| GPM1 (2.80) | RPS19B (−3.17) | YOR293C-A (0.01) |
| MET22 (2.79) | TMA19 (−3.17) | SIN3 (0.01) |
| WTM1 (2.69) | RPL16A (−3.17) | FYV12 (0.01) |
| ARN2 (2.68) | RPL37A (−3.16) | TRP3 (0.01) |
| RTS3 (2.66) | RPL19A (−3.16) | SMK1 (0.01) |
| CPR1 (2.63) | RPL7A (−3.16) | PFD1 (0.01) |
| YER067W (2.62) | RPS17A (−3.15) | YGR146C-A (0.01) |
| GSH1 (2.61) | RPL17B (−3.14) | KAR9 (0.01) |
| PGM2 (2.61) | RPS23B (−3.12) | FIG2 (0.01) |
| GSC2 (2.61) | RPS27B (−3.11) | PGA1 (0.01) |
| PRC1 (2.60) | RPL20B (−3.11) | YKR051W (0.01) |
| MET8 (2.56) | RPS11B (−3.10) | SMC2 (0.01) |
| HOM6 (2.56) | RPL23A (−3.10) | SNF6 (0.01) |
| STF1 (2.55) | RPS28A (−3.08) | YLR412C-A (0.01) |
| GOR1 (2.55) | STM1 (−3.07) | FUS1 (0.01) |
| HNT1 (2.54) | RPL40B (−3.05) | MTW1 (0.01) |
| YET1 (2.52) | RPL34B (−3.05) | CNA1 (0.01) |
| YHR138C (2.52) | RPP1B (−3.04) | CDC12 (0.01) |
| MMP1 (2.51) | RPL39 (−3.03) | MSC7 (0.01) |
| HSP78 (2.51) | ILV3 (−3.02) | SMC6 (0.01) |
| HXK1 (2.51) | RPL23B (−2.99) | YMC1 (0.01) |

Table S2: Lists of 100 most up-regulated, 100 most down-regulated, and 100 most constant genes after 60 minutes of cadmium treatment. Upmost and downmost are sorted with greatest change first (log-fold expression change in parentheses). Most constant are sorted with least absolute change first (absolute log-fold expression change in parentheses).

| RoboCOP | COMPETE | random |
|---|---|---|
| 0.77 | 0.54 | 0.52 |

Table S3: F1-score of nucleosome positions predicted by RoboCOP, COMPETE, and randomly selected nucleosome positions. COMPETE predicted nucleosomes are only slightly better than random prediction of nucleosome positions.

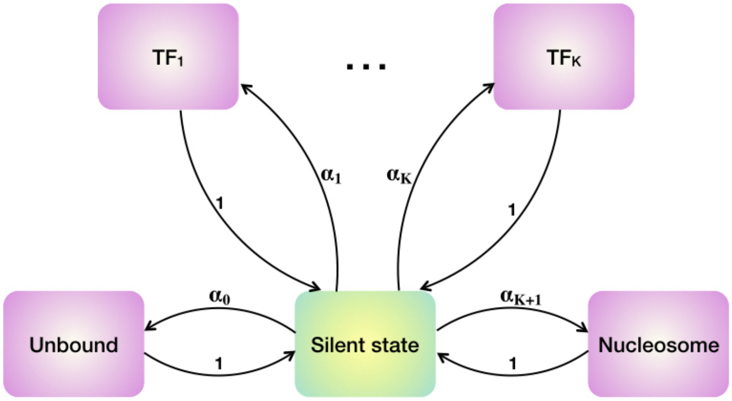


Figure S1: Transition diagram among the different DBFs. The DBFs are nucleosome, $K$ number of TFs and unbound state. The model transitions to any DBF (pink), say $\pi_k$, with probability $\alpha_k$ and then comes back to the silent state (green) with a probability of 1. From the silent state it can then transition to another DBF. The transitions among the DBFs are therefore independent of one another.

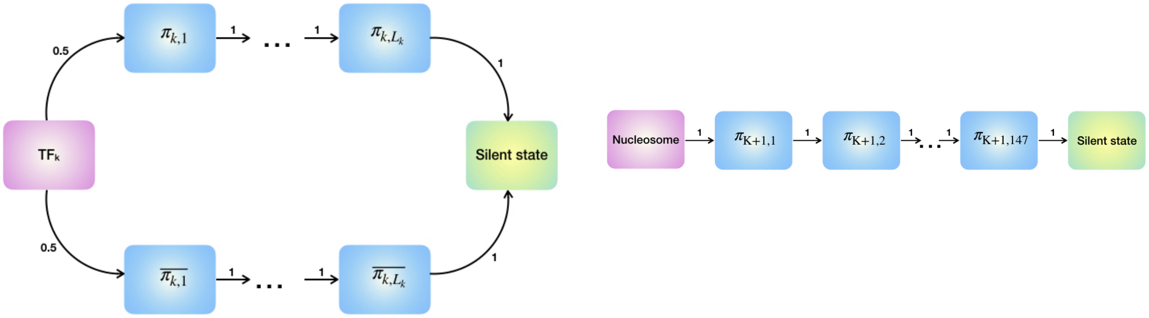


(a) (b)

Figure S2: (a) State transition diagram of a TF. From the first hidden silent state (pink) of the TF, the model can transition to the forward motif (top) or the reverse motif (bottom) with equal probability of 0.5. It then transitions through the entire length, $L_k$, of the motif. After transitioning through the entire motif, the model transitions back to the central silent state (green) with probability of 1. (b) State transition diagram of the nucleosome model, $\pi_{K+1}$. Nucleosomes are modeled to be 147 bases long. After transitioning through the 147 nucleosome positions the model transitions back to the central silent state.

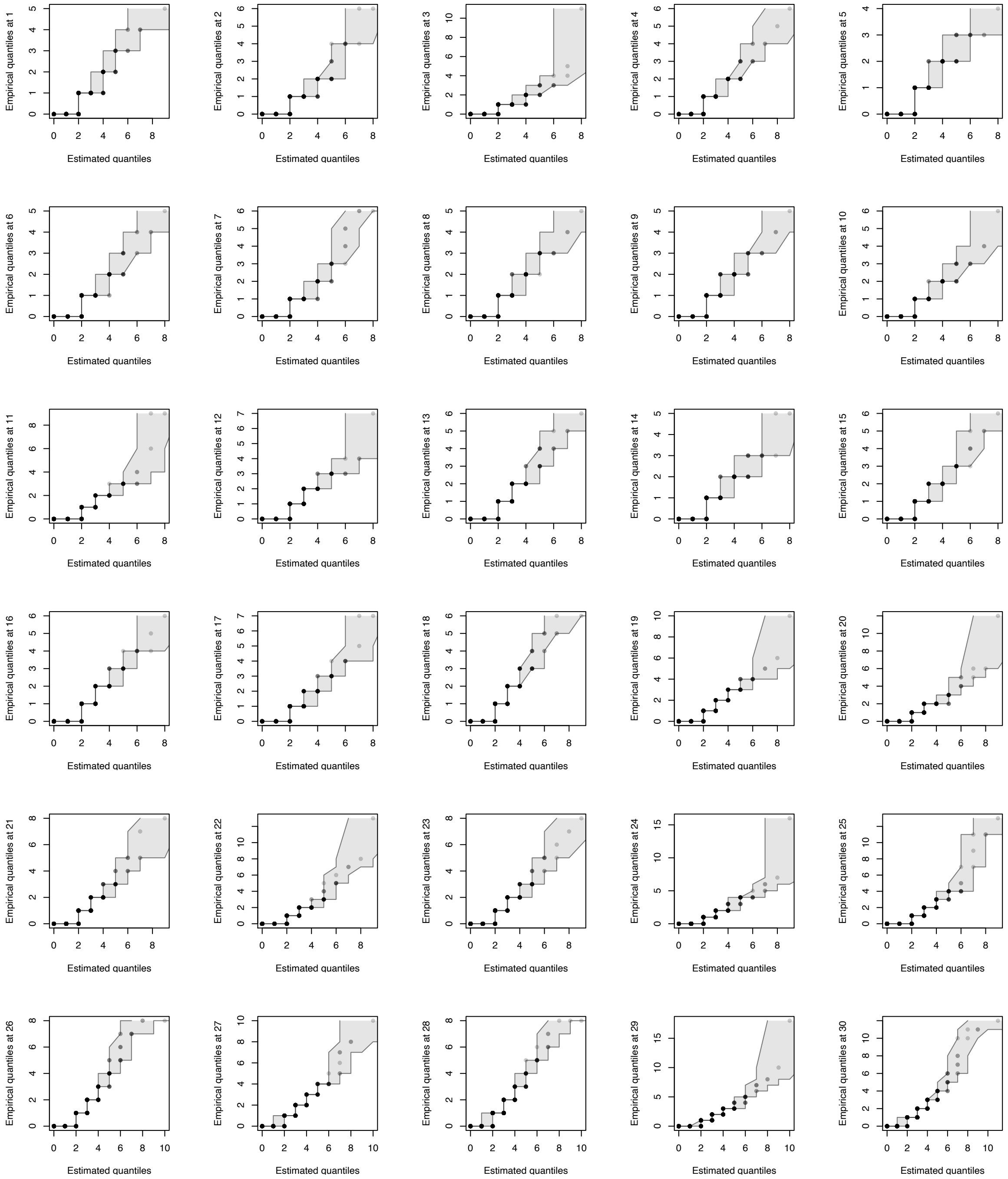
Empirical quantiles at 1
Estimated quantiles
Empirical quantiles at 2
Estimated quantiles
Empirical quantiles at 3
Estimated quantiles
Empirical quantiles at 4
Estimated quantiles
Empirical quantiles at 5
Estimated quantiles
Empirical quantiles at 6
Estimated quantiles
Empirical quantiles at 7
Estimated quantiles
Empirical quantiles at 8
Estimated quantiles
Empirical quantiles at 9
Estimated quantiles
Empirical quantiles at 10
Estimated quantiles
Empirical quantiles at 11
Estimated quantiles
Empirical quantiles at 12
Estimated quantiles
Empirical quantiles at 13
Estimated quantiles
Empirical quantiles at 14
Estimated quantiles
Empirical quantiles at 15
Estimated quantiles
Empirical quantiles at 16
Estimated quantiles
Empirical quantiles at 17
Estimated quantiles
Empirical quantiles at 18
Estimated quantiles
Empirical quantiles at 19
Estimated quantiles
Empirical quantiles at 20
Estimated quantiles
Empirical quantiles at 21
Estimated quantiles
Empirical quantiles at 22
Estimated quantiles
Empirical quantiles at 23
Estimated quantiles
Empirical quantiles at 24
Estimated quantiles
Empirical quantiles at 25
Estimated quantiles
Empirical quantiles at 26
Estimated quantiles
Empirical quantiles at 27
Estimated quantiles
Empirical quantiles at 28
Estimated quantiles
Empirical quantiles at 29
Estimated quantiles
Empirical quantiles at 30
Estimated quantiles

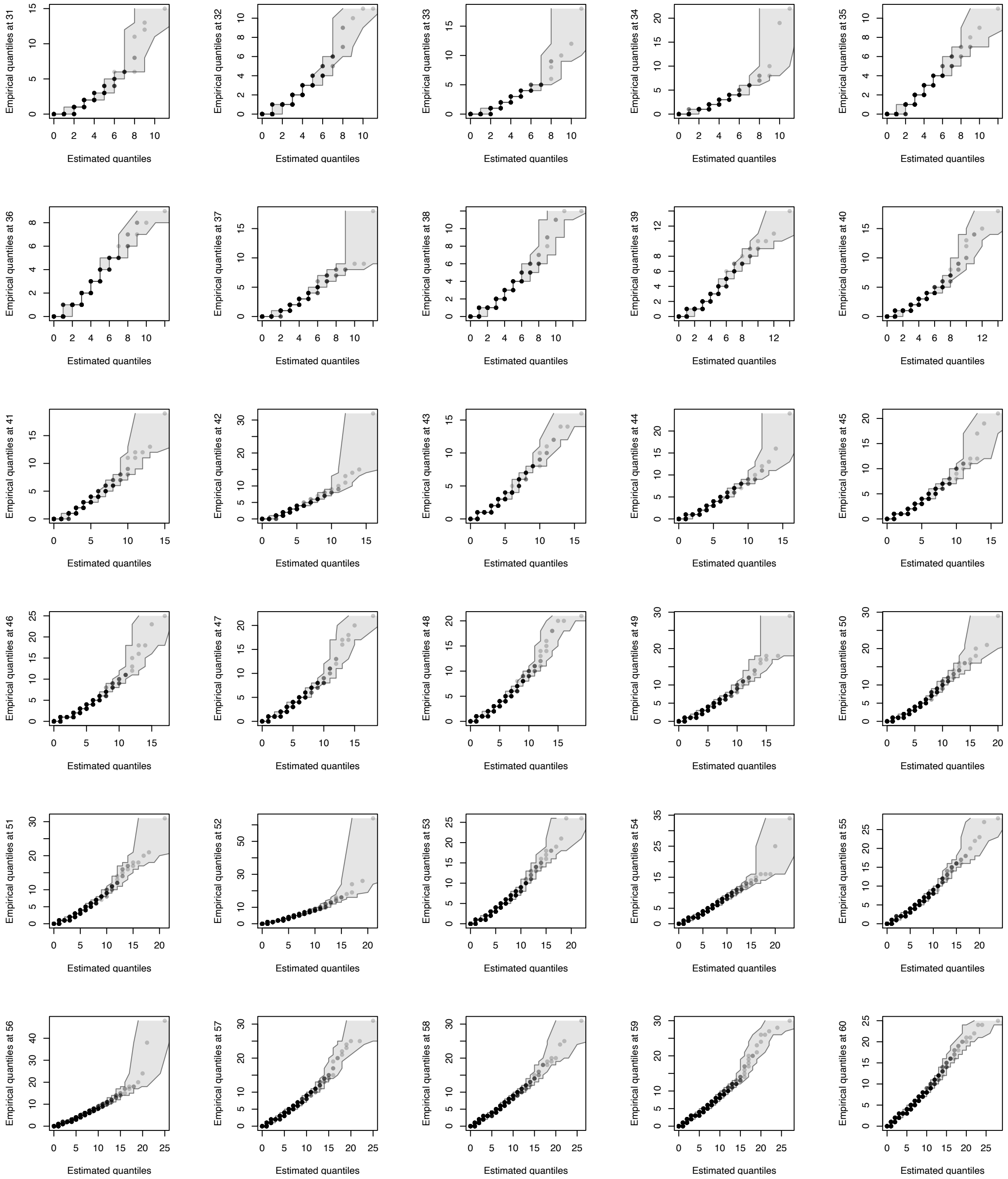
Empirical quantiles at 31
Empirical quantiles at 32
Empirical quantiles at 33
Empirical quantiles at 34
Empirical quantiles at 35
Empirical quantiles at 36
Empirical quantiles at 37
Empirical quantiles at 38
Empirical quantiles at 39
Empirical quantiles at 40
Empirical quantiles at 41
Empirical quantiles at 42
Empirical quantiles at 43
Empirical quantiles at 44
Empirical quantiles at 45
Empirical quantiles at 46
Empirical quantiles at 47
Empirical quantiles at 48
Empirical quantiles at 49
Empirical quantiles at 50
Empirical quantiles at 51
Empirical quantiles at 52
Empirical quantiles at 53
Empirical quantiles at 54
Empirical quantiles at 55
Empirical quantiles at 56
Empirical quantiles at 57
Empirical quantiles at 58
Empirical quantiles at 59
Empirical quantiles at 60
Estimated quantiles

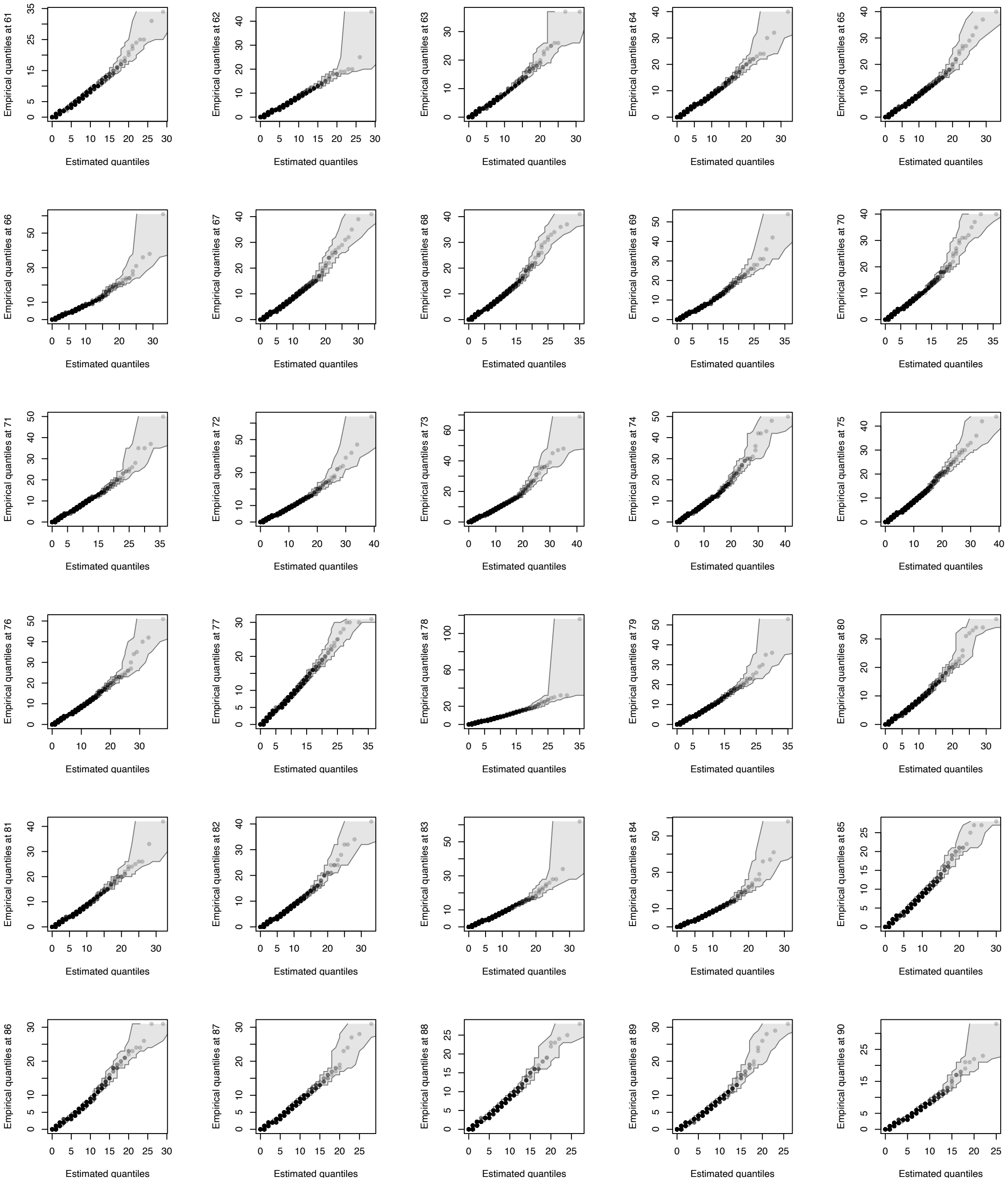
Empirical quantiles at 61
Empirical quantiles at 62
Empirical quantiles at 63
Empirical quantiles at 64
Empirical quantiles at 65
Empirical quantiles at 66
Empirical quantiles at 67
Empirical quantiles at 68
Empirical quantiles at 69
Empirical quantiles at 70
Empirical quantiles at 71
Empirical quantiles at 72
Empirical quantiles at 73
Empirical quantiles at 74
Empirical quantiles at 75
Empirical quantiles at 76
Empirical quantiles at 77
Empirical quantiles at 78
Empirical quantiles at 79
Empirical quantiles at 80
Empirical quantiles at 81
Empirical quantiles at 82
Empirical quantiles at 83
Empirical quantiles at 84
Empirical quantiles at 85
Empirical quantiles at 86
Empirical quantiles at 87
Empirical quantiles at 88
Empirical quantiles at 89
Empirical quantiles at 90
Estimated quantiles

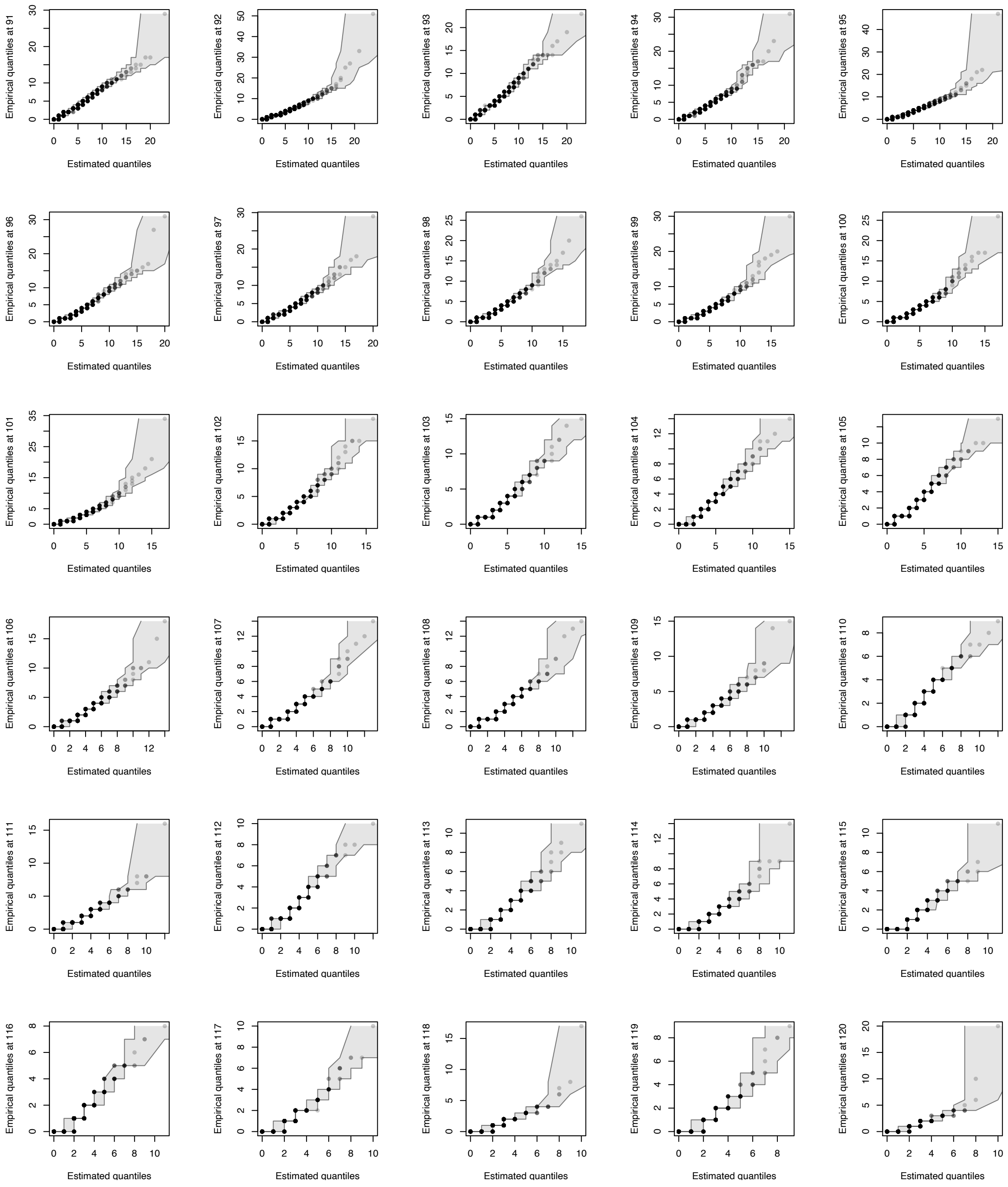
Empirical quantiles at 91
Estimated quantiles
Empirical quantiles at 92
Estimated quantiles
Empirical quantiles at 93
Estimated quantiles
Empirical quantiles at 94
Estimated quantiles
Empirical quantiles at 95
Estimated quantiles
Empirical quantiles at 96
Estimated quantiles
Empirical quantiles at 97
Estimated quantiles
Empirical quantiles at 98
Estimated quantiles
Empirical quantiles at 99
Estimated quantiles
Empirical quantiles at 100
Estimated quantiles
Empirical quantiles at 101
Estimated quantiles
Empirical quantiles at 102
Estimated quantiles
Empirical quantiles at 103
Estimated quantiles
Empirical quantiles at 104
Estimated quantiles
Empirical quantiles at 105
Estimated quantiles
Empirical quantiles at 106
Estimated quantiles
Empirical quantiles at 107
Estimated quantiles
Empirical quantiles at 108
Estimated quantiles
Empirical quantiles at 109
Estimated quantiles
Empirical quantiles at 110
Estimated quantiles
Empirical quantiles at 111
Estimated quantiles
Empirical quantiles at 112
Estimated quantiles
Empirical quantiles at 113
Estimated quantiles
Empirical quantiles at 114
Estimated quantiles
Empirical quantiles at 115
Estimated quantiles
Empirical quantiles at 116
Estimated quantiles
Empirical quantiles at 117
Estimated quantiles
Empirical quantiles at 118
Estimated quantiles
Empirical quantiles at 119
Estimated quantiles
Empirical quantiles at 120
Estimated quantiles

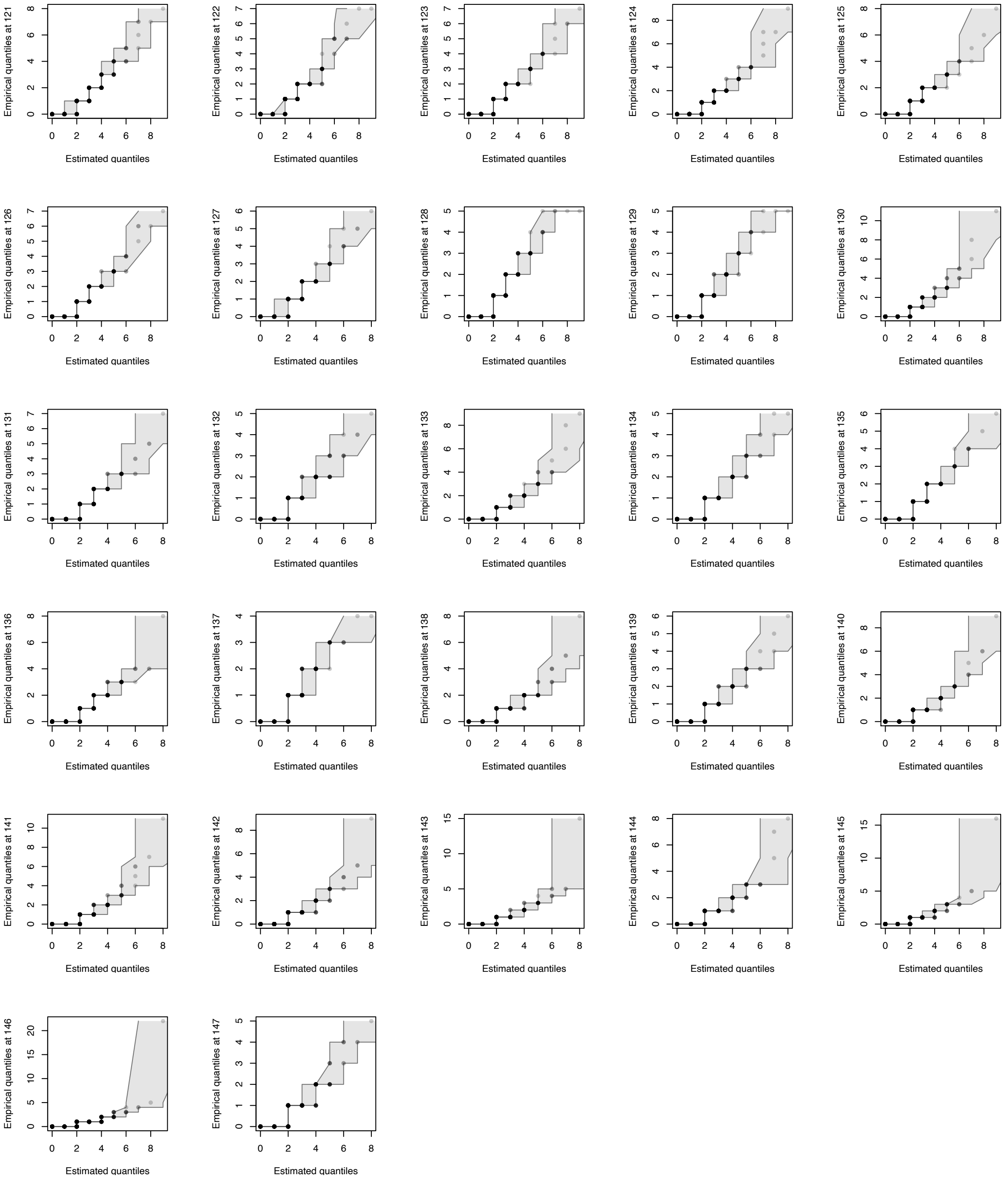


Figure S3: Quantile-quantile plots of negative binomial distributions for `nucFrags` midpoint counts at each of the 147 positions of a nucleosome. The shaded gray region corresponds to the 95% confidence interval of the plots. The plots show the estimated negative binomial distributions are good fits to the midpoint counts of `nucFrags`.

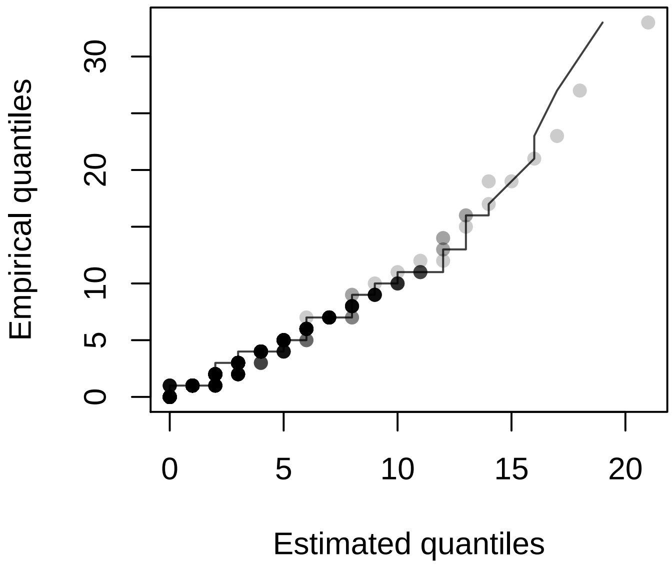


Figure S4: Quantile-quantile plot of negative binomial distribution for `shortFrags` midpoint counts at annotated Abf1 and Reb1 binding sites (1). The plot shows the estimated negative binomial distribution is a good fit to the midpoint counts of `shortFrags`.

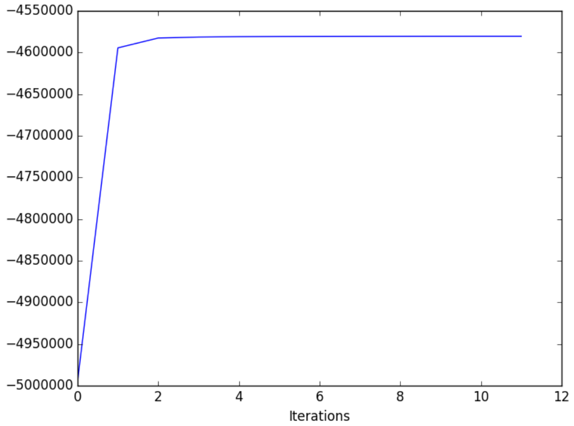


Figure S5: Likelihood in the 10 iterations of EM of RoboCOP. The likelihood converges in 10 iterations.

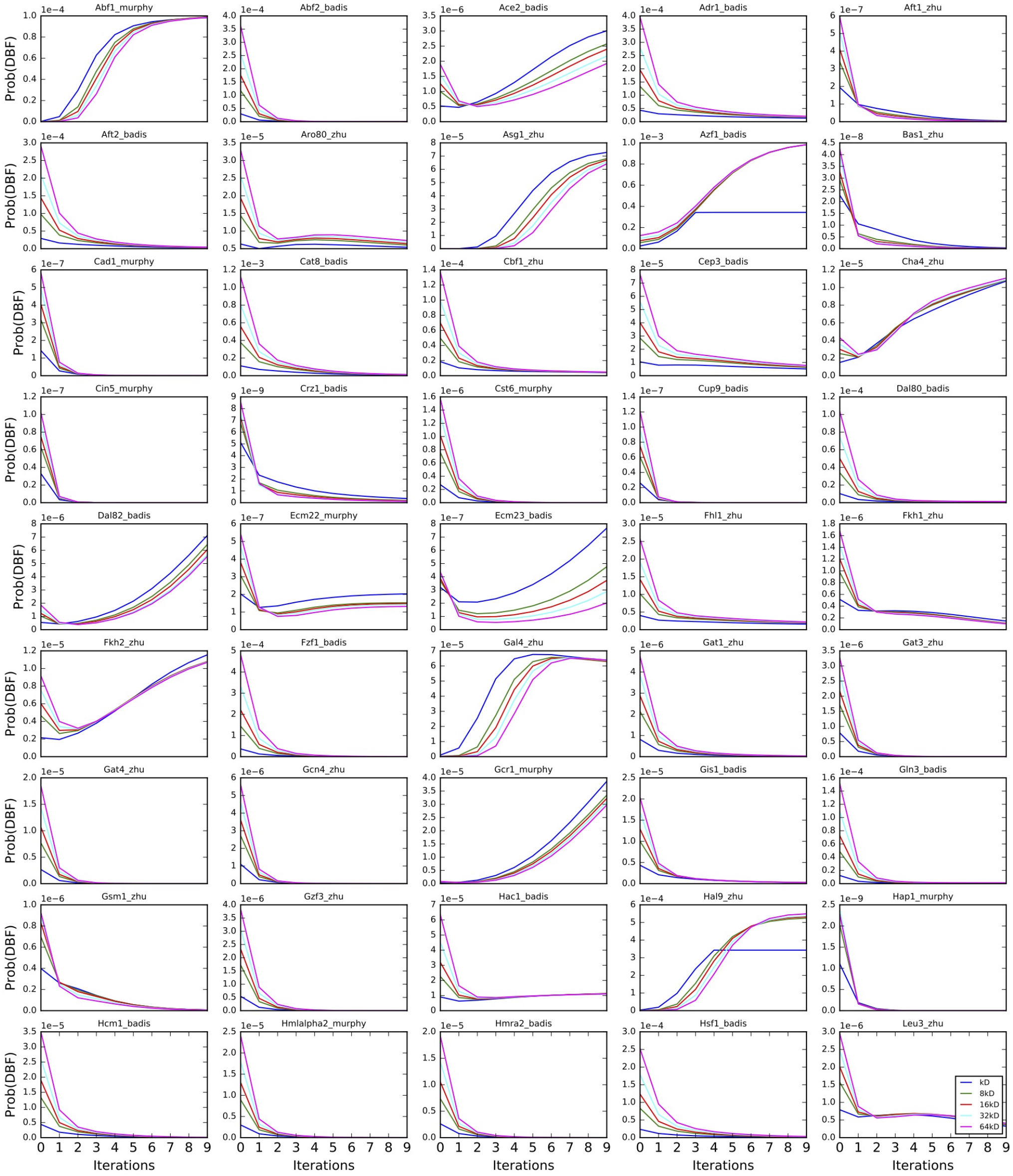
Abf1_murphy
Abf2_badis
Ace2_badis
Adr1_badis
Aft1_zhu
Aft2_badis
Aro80_zhu
Asg1_zhu
Azf1_badis
Bas1_zhu
Cad1_murphy
Cat8_badis
Cbf1_zhu
Cep3_badis
Cha4_zhu
Cin5_murphy
Crz1_badis
Cst6_murphy
Cup9_badis
Dal80_badis
Dal82_badis
Ecm22_murphy
Ecm23_badis
Fhl1_zhu
Fkh1_zhu
Fkh2_zhu
Fzf1_badis
Gal4_zhu
Gat1_zhu
Gat3_zhu
Gat4_zhu
Gcn4_zhu
Gcr1_murphy
Gis1_badis
Gln3_badis
Gsm1_zhu
Gzf3_zhu
Hac1_badis
Hal9_zhu
Hap1_murphy
Hcm1_badis
Hmlalpha2_murphy
Hmra2_badis
Hsf1_badis
Leu3_zhu
Prob(DBF)
Iterations
kD
8kD
16kD
32kD
64kD

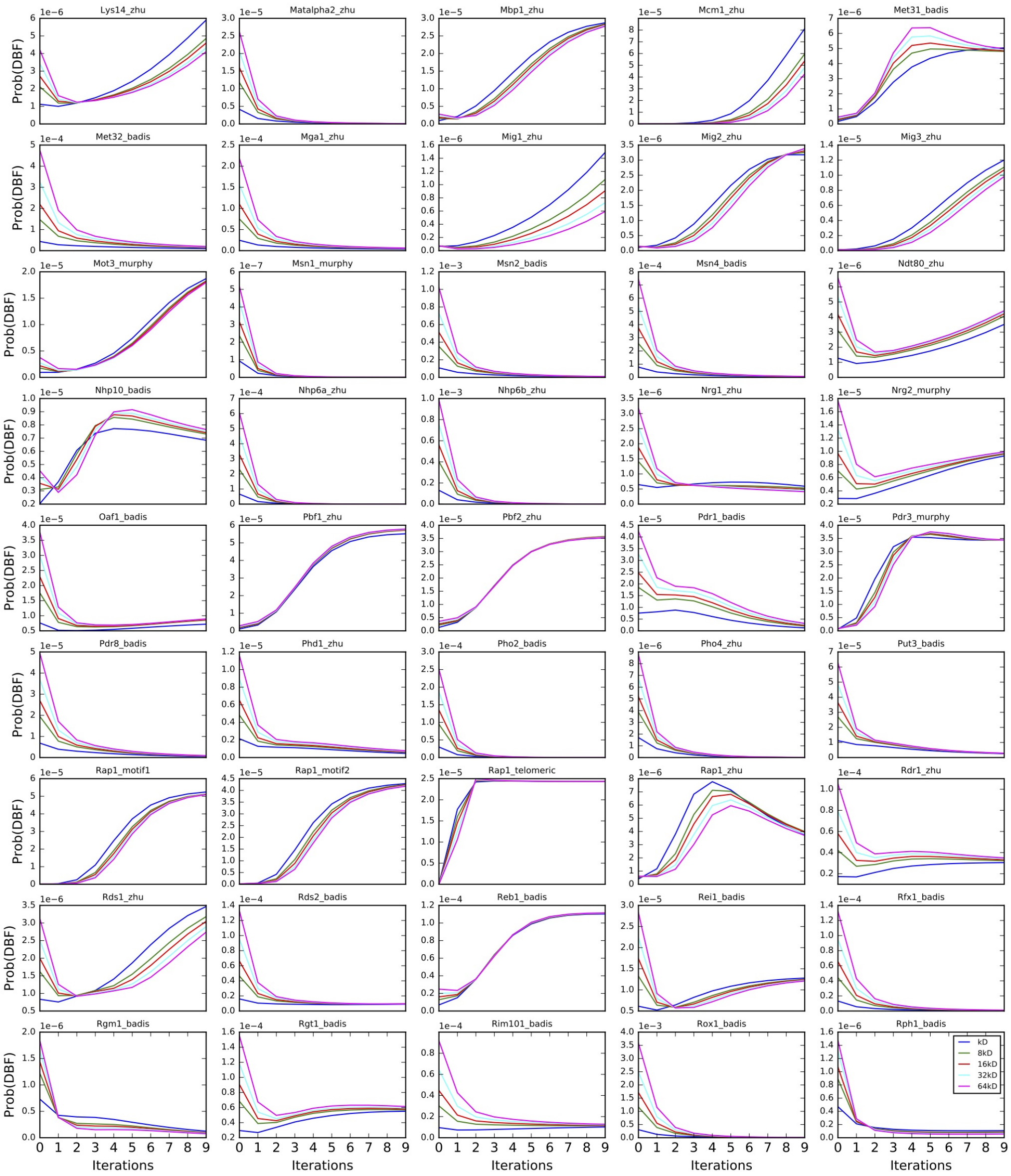
Lys14_zhu
Matalpha2_zhu
Mbp1_zhu
Mcm1_zhu
Met31_badis
Met32_badis
Mga1_zhu
Mig1_zhu
Mig2_zhu
Mig3_zhu
Mot3_murphy
Msn1_murphy
Msn2_badis
Msn4_badis
Ndt80_zhu
Nhp10_badis
Nhp6a_zhu
Nhp6b_zhu
Nrg1_zhu
Nrg2_murphy
Oaf1_badis
Pbf1_zhu
Pbf2_zhu
Pdr1_badis
Pdr3_murphy
Pdr8_badis
Phd1_zhu
Pho2_badis
Pho4_zhu
Put3_badis
Rap1_motif1
Rap1_motif2
Rap1_telomeric
Rap1_zhu
Rdr1_zhu
Rds1_zhu
Rds2_badis
Reb1_badis
Rei1_badis
Rfx1_badis
Rgm1_badis
Rgt1_badis
Rim101_badis
Rox1_badis
Rph1_badis
Prob(DBF)
Iterations
kD
8kD
16kD
32kD
64kD

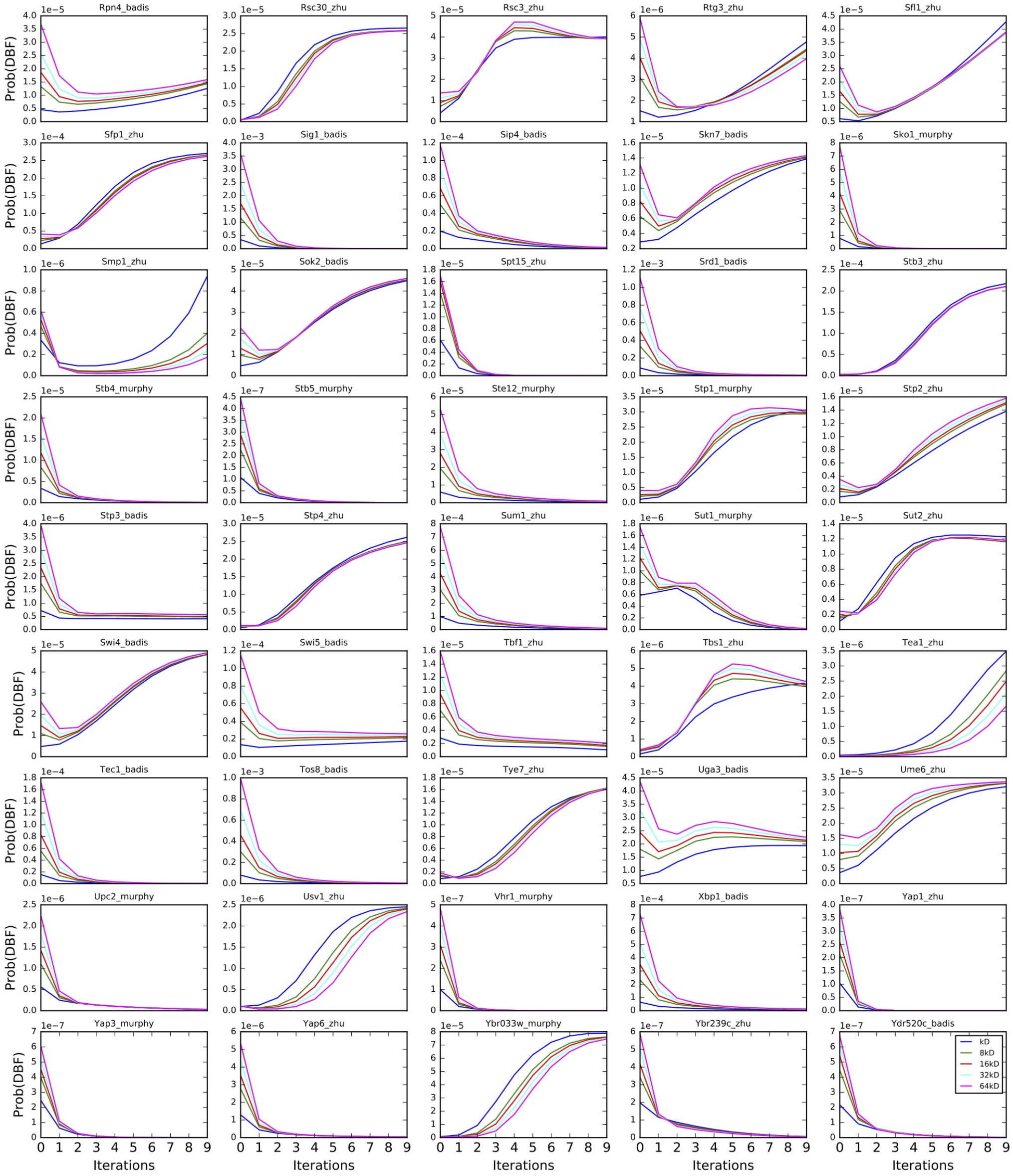
Rpn4_badis
Rsc30_zhu
Rsc3_zhu
Rtg3_zhu
Sfl1_zhu
Sfp1_zhu
Sig1_badis
Sip4_badis
Skn7_badis
Sko1_murphy
Smp1_zhu
Sok2_badis
Spt15_zhu
Srd1_badis
Stb3_zhu
Stb4_murphy
Stb5_murphy
Ste12_murphy
Stp1_murphy
Stp2_zhu
Stp3_badis
Stp4_zhu
Sum1_zhu
Sut1_murphy
Sut2_zhu
Swi4_badis
Swi5_badis
Tbf1_zhu
Tbs1_zhu
Tea1_zhu
Tec1_badis
Tos8_badis
Tye7_zhu
Uga3_badis
Ume6_zhu
Upc2_murphy
Usv1_zhu
Vhr1_murphy
Xbp1_badis
Yap1_zhu
Yap3_murphy
Yap6_zhu
Ybr033w_murphy
Ybr239c_zhu
Ydr520c_badis
Prob(DBF)
Iterations
kD
8kD
16kD
32kD
64kD

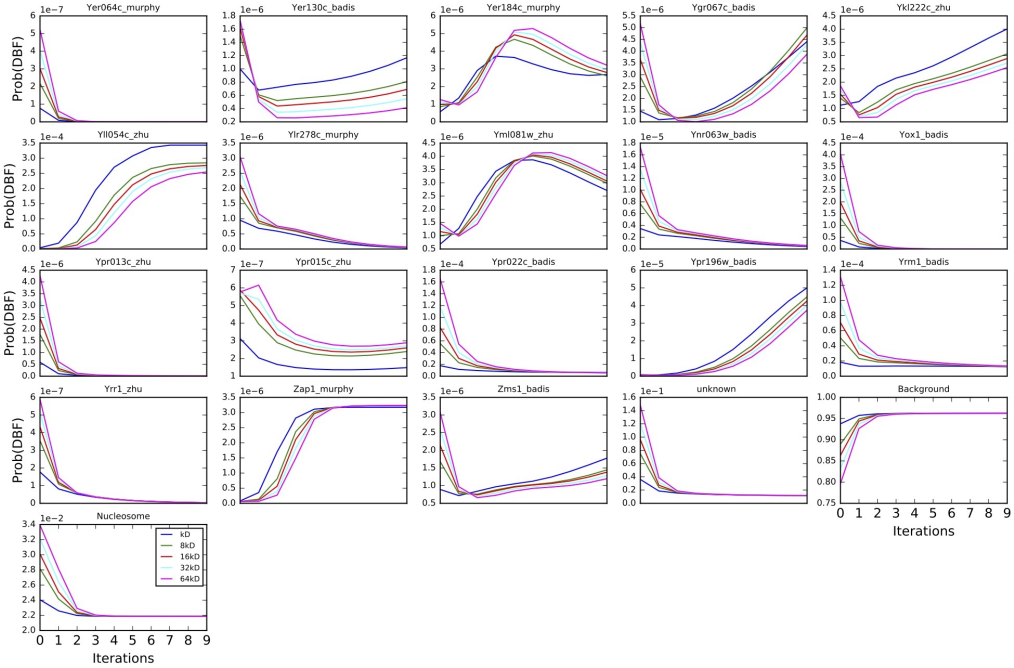


Figure S6: Transition weights of DBFs in every iteration of EM when initializing using multiples of $K_D$.

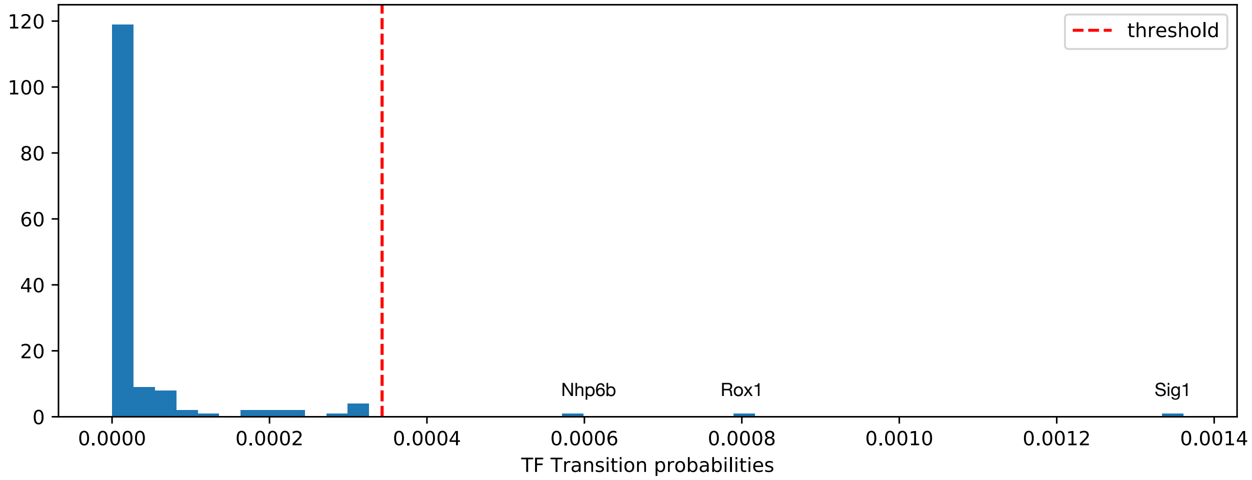


Figure S7: Choosing threshold (dotted red line) for transition probabilities of TFs. Threshold value is calculated to be 2 standard deviations more than the mean of all initial TF transition probabilities. Only three TFs (Nhp6b, Rox1, and Sig1) have an initial transition probability greater than the threshold.

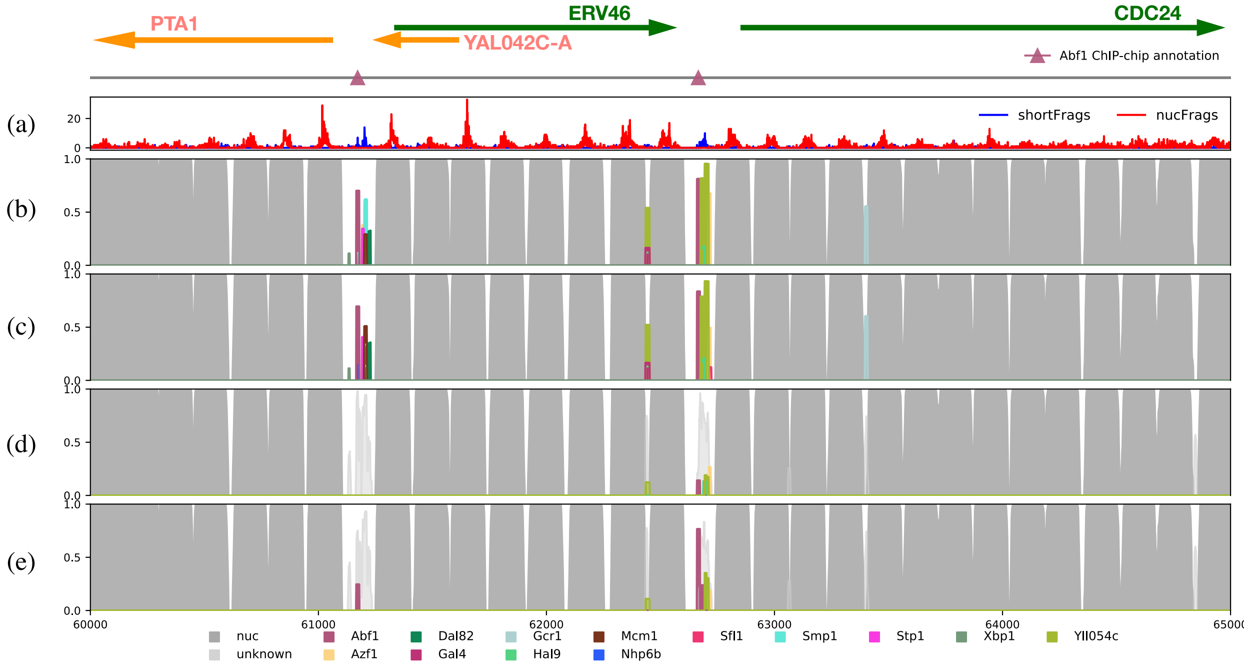


Figure S8: (a) Aggregate of MNase-seq `nucFrags` and `shortFrags` from positions 60,000 to 65,000 of yeast chromosome I. Above the plot are genomic annotations for this region, with Watson strand genes depicted as green arrows and Crick strand genes as orange. Below the gene annotations, known TF binding sites (1) are indicated using triangles. This region contains two annotated binding sites for Abf1 (pink). (b) Running RoboCOP without a threshold and without an 'unknown' factor, (c) with a threshold but without an 'unknown' factor, (d) without a threshold but with an 'unknown' factor, and (e) with a threshold and with an 'unknown' factor (i.e., the final model). In (b,c), we observe large TF clusters that result in increased false positive predictions. In (d), without the threshold, we see that most TFs are predicted as being an 'unknown' factor. In (e), by adding an 'unknown' factor and a threshold on how large a TF transition probability can grow, we achieve a happy medium.

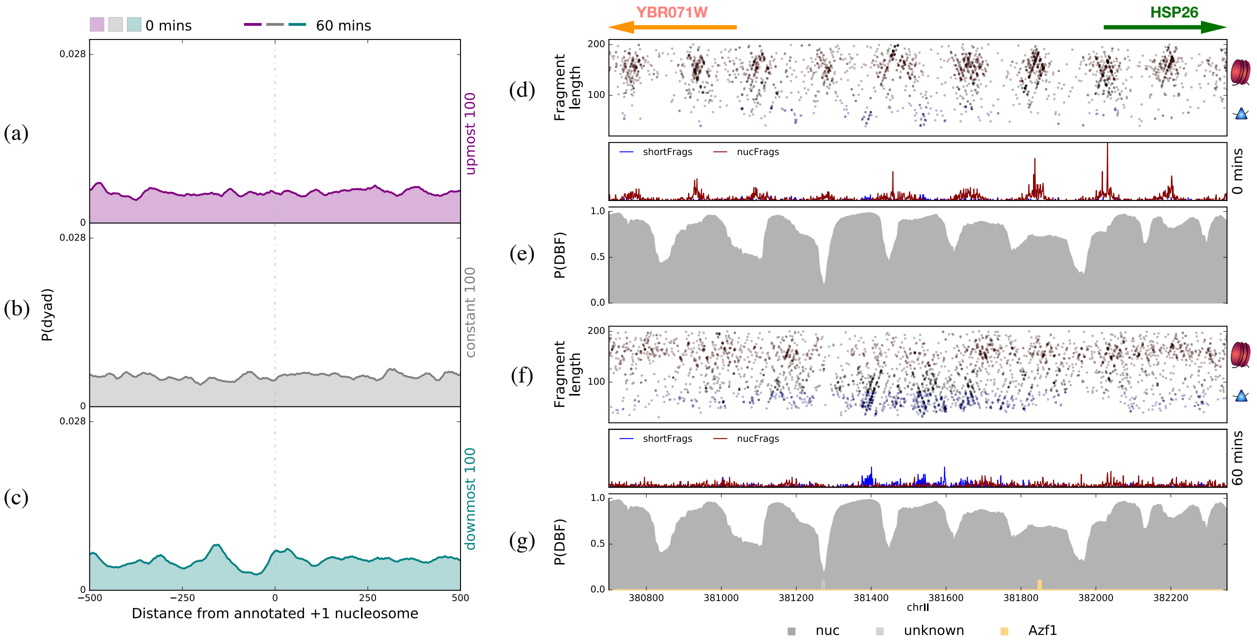


Figure S9: (a–c) Aggregate nucleosome dyad probability, as computed by COMPETE, around annotated +1 nucleosomes (5) of (a) the 100 most up-regulated genes (purple), (b) the 100 genes least changed in transcription (gray), and (c) the 100 most down-regulated genes (teal), before and 60 minutes after treating cells with cadmium. We find that COMPETE does not identify any change in these nucleosome occupancy profiles after cadmium treatment. (d) Two-dimensional plot of MNase-seq fragments near the HSP26 promoter (positions 380,700 to 382,350 of yeast chromosome II are shown) before treatment with cadmium (`nucFrags` in red; `shortFrags` in blue), along with the `nucFrags` and `shortFrags` signals that result from aggregating those midpoint counts. Gene annotations depicted with arrows at the top (Watson strand in green; Crick strand in orange). (e) COMPETE-predicted occupancy profile of this region before treatment with cadmium. (f,g) The same as (d,e), respectively, but 60 minutes after cadmium treatment. Even though the MNase-seq data have changed significantly after treatment (d,f), COMPETE does not make use of this data, so fails to capture the changes to the chromatin occupancy profile of the genome.
